# ICOS expression is required for maintenance but not the formation of germinal centers in the spleen in response to *P. yoelii* infection

**DOI:** 10.1101/2021.08.24.457597

**Authors:** Kara A. O’Neal, Leah E. Latham, Enatha Ntirandekura, Camille L. Foscue, Jason S. Stumhofer

**Author notes:** These authors contributed equally to the work (Kara A. O’Neal, Leah E. Latham). Correspondence should be addressed to J.S.S., University of Arkansas for Medical Sciences, Department of Microbiology and Immunology, Room 521A, 4301 W. Markham St., Little Rock, AR 72205.

## Abstract

Inducible T cell co-stimulator (ICOS) plays a key role in the differentiation and maintenance of follicular helper T (Tfh) cells and thus germinal center (GC) formation. Previously, our lab showed in a *Plasmodium chabaudi* infection model that *Icos*^-/-^ mice did not form GCs despite a persistent infection and thus a continued antigen (Ag) load. Here, we show that resolution of a primary infection with *P. yoelii*, was delayed in *Icos*^-/-^ mice. This phenotype was associated with a reduction in the accumulation of Tfh-like and GC Tfh cells and an early deficiency in Ag-specific antibody (Ab) production. However, *Icos*^-/-^ mice maintained their ability to form GCs, though they were less frequent in number than in wild-type (WT) mice. Furthermore, while Ab production in *Icos^-/-^* mice matched that of WT mice after the infection cleared, the Abs lacked signs of affinity maturation, suggesting functional defects associated with these GCs. Eventually, these GC structures dissipated more rapidly in *Icos^-/-^* mice than in WT mice. Moreover, the ability of *Icos^-/-^* mice to form these GC structures is not reliant on the high Ag load associated with *P. yoelii* infections, as GC formation was preserved in *Icos^-/-^* mice treated with early with atovaquone. Finally, mice were unable to form secondary GCs in the absence of ICOS after re-challenge. Overall, these data demonstrate the necessity of ICOS in the maintenance of Tfh cells, the formation and maintenance of sufficient numbers of functioning GCs, and the ability to generate new GC structures after re-infection with *P. yoelii*.

## Introduction

It is well established that antibody (Ab) production is crucial for controlling a *Plasmodium* infection [1]. Germinal centers (GC), secondary follicles that develop within existing B cell follicles, are essential for generating high affinity Abs through the process of somatic hypermutation. Those GC B cells that develop favorable mutations in their B-cell receptor that improve their affinity for binding Ag eventually differentiate into long-lived memory B cells (MBCs) and plasma cells. Within the GC, B cells rely on CD4^+^ T cell help from an effector population known as follicular helper T (Tfh) cells. During *Plasmodium* infection in mice, CD4^+^ T cell differentiation yields two predominant populations of Tfh cells: Tfh-like cells (CXCR5^+^PD-1^+^CXCR3^+^Bcl6^+^Tbet^+^) that display a mixed Tfh-Th1 phenotype and GC Tfh cells (CXCR5^+^PD-1^++^CXCR3^-^Bcl6^+^Tbet^lo/-^) that localize to the GC [2]. Tfh cells provide signals that promote the proliferation, survival, and differentiation of B cells before and during their time within the GC. Likewise, B cells reciprocate help to T cells via engagement of various co-stimulatory molecules. One of these co-stimulatory molecules, ICOS (inducible co-stimulator molecule), interacts with its ligand (ICOSL) on B cells to promote Tfh cell entry into the B cell follicle and the perpetuation of the Tfh cell phenotype [3]. In the absence of ICOS-ICOSL interactions, Tfh cell development and GC formation are impaired, leading to an Ab response dominated by low-affinity Ab production [4].

Previously, our lab showed that *P. chabaudi* infection in *Icos^-/-^* mice resulted in a reduced acute parasite burden due to an enhanced Th1 response. While class-switched parasite-specific Ab production occurred, GC formation was absent, which led to the inability of *Icos^-/-^* mice to resolve their persistent infection due to a failure to maintain and generate high-affinity Ab titers. The absence of GC formation coincided with the inability of the mice to maintain Tfh cell numbers in the spleen and the failure to generate GC Tfh cells [5]. Here, we are interested in determining if the role of ICOS mirrors that observed during *P. chabaudi* infection in mice infected with another non-lethal *Plasmodium* species, *P. yoelii*, in which the humoral response is critical for control of the primary infection [6]. Here, *Icos^-/-^* mice infected with *P. yoelii* displayed an exacerbated peak parasitemia and delayed resolution of parasite clearance. While Tfh-like and GC Tfh cell accumulation and maintenance were impaired in the absence of ICOS signaling, surprisingly, GC structures were apparent in the spleen after the resolution of the infection. However, the GC structures deteriorated faster in *Icos^-/-^* mice than wild-type (WT) mice.

While there is a consensus understanding that Tfh cell maturation, migration, and function are dependent on cognate Ag and ICOS-L delivery by B cells [8, 9], the requirement of ICOS-L in these events is suggested as being only operative in settings of limited Ag delivery by cognate B cells [10, 11]. Hence, when Ag is abundant, any antigen-presenting cell is equipped and capable of directing Tfh cell differentiation. Therefore, based on these findings, we hypothesized that the high parasite burden associated with *P. yoelii* infection contributed to the ability of these mice to form GCs in the absence of ICOS-L:ICOS signaling. However, here we show that treatment with the anti-malarial drug atovaquone (AV) during the early stages of the infection does not restrict the ability of *Icos^-/-^* mice to produce GCs, indicating that a high Ag load is not the sole factor allowing for the formation of GCs in the absence of ICOS.

## Materials and Methods

### Mice and infections

Male and female WT (C57BL/6J) and *Icos*^-/-^ (B6.129P2-Icostm1Mak/J) were purchased from The Jackson Laboratory. Balb/c mice were purchased from Charles River. In compliance with institutional guidelines, mice were bred and housed in specific-pathogen-free facilities at the University of Arkansas for Medical Sciences. The IACUC at the University of Arkansas for Medical Sciences approved all procedures on mice in this study. Studies were performed in male and female mice to account for sex differences in the immune response. For *Plasmodium yoelii* infections, male BALB/c mice were injected intraperitoneally (i.p.) with cryopreserved parasitized red blood cell (pRBC) stocks (MR4 17X, BEI Resources Repository). From each passage, 10^5^ pRBCs were injected i.p. for infections in experimental mice. Mice were infected between seven and ten weeks of age. Flow cytometry was used to determine blood parasitemia based on a previously described method. All procedures involving *P. yoelii* 17X were approved by the IBC at the University of Arkansas for Medical Sciences.

### Flow cytometry and antibodies

Total splenocytes were processed and filtered through a 40 μm cell strainer and subjected to RBC lysis (0.86% NH4Cl solution) to create single cell suspensions for flow cytometry analysis. Throughout the process, cells were kept in complete RPMI (RPMI 1640, 10% fetal plex, 10% non-essential amino acids, 10% sodium pyruvate, 10% L-glutamine, 10% penicillin and streptomycin, and 1% 2-βME). To prevent non-specific Ab binding, Fc receptors on cells were blocked with anti-mouse CD16/32 (24G2; BioXCell) in a buffer containing normal mouse and rat IgG (Life Technologies). FACS buffer (1× PBS, 0.2% BSA and 0.2% 0.5M EDTA), was used for washing cells. Ab cocktails were prepared in FACs buffer supplemented with Super Bright Staining Buffer (ThermoFisher) for surface staining. Upon completion of surface staining, cells were fixed in 4% paraformaldehyde (Electron Microscopy Sciences) if intracellular staining was not required. Fluorescence minus one (FMO) controls were used to set positive gates. An LSRIIFortessa (Becton Dickson) was used to acquire samples. Data were analyzed using FlowJo version 10 software.

### Intracellular staining

For intracellular staining of transcription factors, a Foxp3/Transcription factor staining buffer set (eBioscience) was used per the manufacturer’s direction. For cytokine staining, processed splenocytes, prior to surface staining, were incubated with PMA and ionomycin in the presence of Brefeldin A (Sigma) at 37°C for 4 hours. After incubation, cells underwent surface staining followed by fixation with 4% PFA and permeabilization with 0.1% saponin diluted in FACS buffer. Cells were then subjected to cytokine staining with IFN-γ, IL-10, IL-21, and TNF-α. To detect IL-21, cells were first incubated with recombinant murine IL-21 receptor (R&D systems) then incubated with human Fc receptor coupled to phycoerythrin (ThermoFisher). To set positive gates, fluorescence minus one (FMO) controls were used. Antibodies used for flow cytometry are listed in Supplementary Table 1.

### Immunofluorescence

Splenic sections were embedded in OCT, subjected to flash freezing in liquid nitrogen, and then cut into 4 μm sections. For staining, slides were fixed for 10 minutes at -20°C in an ice-cold 25% acetone/75% methanol solution. Following fixation, slides were blocked with 2% normal goat serum and then stained with CD3ε, GL-7, and IgD (ThermoFisher) at 4°C overnight. Secondary staining consisted of goat anti-rat IgG (H+L)-AF647 and goat anti-Armenian hamster IgG (H+L)-Cy3 Abs at room temperature for 1 hour followed by staining with a goat anti-rat IgM (μ chain) Ab conjugated to FITC (Jackson Immunoresearch) with the same conditions. Fluoroshield (Electron Microscopy Sciences) was used to mount coverslips. An EVOS FL Auto 2 was used to scan slides, and stitched images were analyzed with ImageJ (NIH).

### ELISAs

Recombinant *P. yoelii* AMA-1 or MSP-119 proteins in sodium bicarbonate buffer were used to coat high binding Immunlon HBX plates (Thermo Scientific) at 4°C overnight. Plates were blocked with 5% Fetal Bovine Serum (FBS; Life Technologies) in PBS. Serum was diluted at 1:50 initially and then serially diluted at 1:3 down the plate. HRP-conjugated IgM or IgG (Southern Biotech) was incubated at 37 °C for 1 hour. For detection, SureBlue substrate (KPL) was used, and stop solution (KPL) was used to halt the reaction. Plates were read at an absorbance of 450 nm on a FLUOStar Omega plate reader (BMG Labtech).

### ELISPOTs

ELISPOT plates (Millipore) were activated with 35% ethanol, washed with sterile PBS, and then coated with recombinant *P. yoelii* AMA-1 or MSP-1_19_ protein and incubated at 4°C overnight.

Plates were washed, and approximately 10^5^ splenocytes were plated and incubated at 37°C overnight. Following overnight incubation, cells were dumped and washed off the plate with 1x PBS and blocked with 5% FBS (Life Technologies) in PBS. APS conjugated IgM and IgG (Southern Biotech) were added and incubated for 1 hour at room temperature. Following incubation, an NBT and BCIP solution (Sigma) was used for the detection of spots. An AID ELISPOT reader was used to read plates, and data was analyzed using AID ELISPOT 7.0 software (AID GmbH).

### Atovaquone Treatment

AV (Sigma) was used to clear parasites from infected mice. WT and *Icos^-/-^* mice were administered AV resuspended in a solution of 40% DMSO in sterile H_2_O at a dose of 14 mg/kg i.p. once a day from days 7-12 of the infection.

### α-IcosL blocking

α-IcosL Ab (clone HK5.3; BioXCell) was used to block ICOS-ICOSL signaling. WT mice were administered 200μg of α-IcosL in sterile PBS i.p. every three days starting one day before infection and continuing throughout infection until the day of sacrifice. α-Rat IgG (Sigma) was administered to control WT mice in the same manner.

### Real-time quantitative PCR

CD4^+^ T cells were enriched from spleens of mice infected for 11 or 36 days with *P. yoelii* 17X, using anti-CD4 microbeads (Miltenyi Biotec) and an AutoMACS Pro cell separator (Miltenyi Biotech). Enriched T cells were stained with CD4, CD44, CD11a, PD-1, and CXCR5 to identify antigen-specific activated Tfh-like and GC Tfh cells. Cells were sorted using a BD FACSAriaIII. Cells were washed with sterile PBS and lysed in Trizol (ThermoFisher). RNA was isolated, and DNAse treated per the manufacturer’s instructions (Zymo Research). Superscript III (Invitrogen) was used for the preparation of complementary DNA. SYBR Green PCR Master Mix (BioRad), primers listed in Supplementary Table II, and a QuantStudio 6 Flex real-time PCR system (Life Technologies) were used to perform real-time quantitative PCR. Data were normalized to the *hprt* housekeeping gene, and the 2^-DCt^ method was used to calculate fold changes.

### Statistics

GraphPad Prism 9 (GraphPad Software, Inc., San Diego, CA) and (R 3.4.3, The R Foundation) were used to perform statistical analyses. The figure legends provide details about specific tests of statistical analysis.

## Results

### The absence of ICOS does not impair the Th1 response

To determine how the loss of ICOS signaling impacts the immune response to *P. yoelii*, WT and *Icos^-/-^* mice were infected with a non-lethal strain of *P. yoelii* (17X), and the resulting parasitemia was monitored. Compared to WT mice, *Icos^-/-^* mice consistently had a higher peak parasite burden and showed a significant delay in resolving the infection, though they eventually cleared the infection (Fig. 1A). Thus, ICOS is required for proper control of infection with *P. yoelii*.

**FIG. 1.**
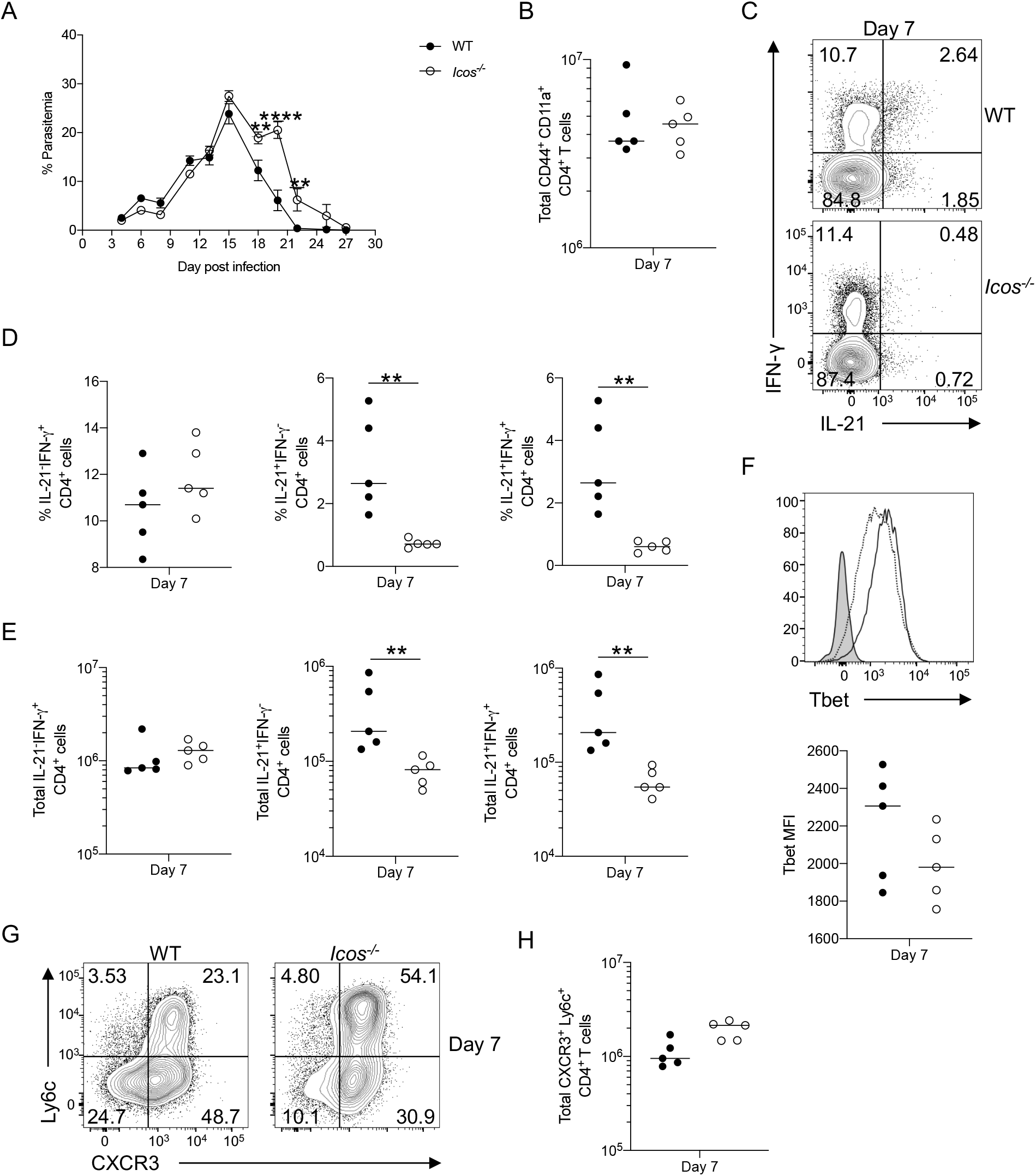
Absence of ICOS does not impair the Th1 response. WT and *Icos^-/-^* mice were injected i.p. with 10^5^ *P. yoelii* pRBCs. (**A**) Representative parasitemia curve during primary infection. (**B**) Total number of live activated (CD44^hi^CD11a^+^) CD4^+^ T cells. (**C**) Representative flow plots of IL-21 and IFN-*γ* expression from live CD4^+^ T cells at day 7 p.i. following *ex vivo* stimulation with PMA and ionomycin in the presence of Brefeldin A. (**D**) Frequency and (**E**) the total number of IL-21^-^IFN-*γ*^+^, IL-21^+^IFN-*γ*^-^, and IL-21^+^IFN-*γ*^+^ CD4^+^ T cells at day 7 p.i. (**F**) Histogram and MFI (median fluorescence intensity) of Tbet expression in live CD44^hi^CD11a^+^ CD4^+^ T cells at day 7 p.i. FMO (fluorescence minus one) is indicated by the shaded peak, WT by a solid line, and *Icos^-/-^* by a dotted line. (**G**) Representative flow plots of Ly6c^+^ and CXCR3^+^ expression on live activated CD4^+^ T cells at day 7 p.i. (**H**) Total number of live activated Ly6c^+^CXCR3^+^ CD4^+^ T cells at day 7 p.i. Data are representative of 3-4 independent experiments with five mice per group. A nonparametric Mann-Whitney *t*-test determined significance. \*\**p* <0.01, \*\*\*\**p* < 0.0001.

Given the described role of ICOS in promoting T and B cell responses [12], we sought to determine how the absence of ICOS impacted these cells during infection with *P. yoelii*. No difference in the activation of antigen-experienced CD4^+^ T cells (CD44^hi^CD11a^+^) was seen in the spleen at day 7 post infection (p.i.) between WT and *Icos^-/-^* mice (Fig. 1B and Supplemental Fig. 1A). CD4^+^ T cells produce several cytokines during acute infection with *P. yoelii*, including IFN-*γ*, TNF*α*, and IL-21 indicative of a mixed Th1 and Tfh cell phenotype [13]. We previously showed that infection with *P. chabaudi* led to impaired IFN-*γ* production by *Icos^-/-^* CD4^+^ T cells [5]. However, a similar defect was not observed in response to *P. yoelii* infection on day 7 p.i. (Fig. 1C-E), or at other times after infection (Supplemental Fig. 1B,C). Alternatively, a significant decrease in both frequency and the total number of IL-21–secreting CD4^+^ T cells, including those co-expressing IFN-*γ*, was seen at day 7 p.i. in *Icos^-/-^* mice (Fig. 1D,E). Although, this decrease in IL-21 production was not sustained as the infection progressed (Supplemental Fig. 1B,C).

During *Plasmodium* infection, two distinct effector phenotypes emerge following activation of CD4^+^ T cells – a Th1 or Tfh phenotype that contribute to parasite clearance [2]. In addition to secreting IFN-*γ*, Th1 cells are characterized by their expression of T-bet, Ly6C, and CXCR3 [14]. Examination of CD4^+^ T cells for the expression of these Th1 associated markers indicated that *Icos^-/-^* CD4^+^ T cells expressed T-bet and the per cell expression of T-bet (median fluorescent intensity [MFI]) was similar between WT and *Icos^-/-^* CD4^+^ T cells at day 7 p.i. (Fig. 1F). Though the frequency of Ly6C^+^CXCR3^+^ CD4^+^ T cells was significantly higher in the *Icos*^-/-^ mice, their total numbers were not very different compared to WT mice (Fig. 1G,H). On day 11 p.i. the CD4^+^ T cells from *Icos^-/-^* mice expressed more T-bet on a per cell basis, and a higher frequency were Ly6C^+^CXCR3^+^, but no difference in the overall number of Ly6C^+^CXCR3^+^ CD4^+^ T cells was seen compared to WT mice (Supplemental Fig. 1D,E). Overall, the data demonstrate that ICOS is unnecessary for T cell activation and the generation and maintenance of a Th1 response during blood-stage *P. yoelii* infection. Still, its loss impacted IL-21 production by CD4^+^ T cells during the early stages of acute infection.

### *Icos^-/-^* mice exhibit defects in Tfh-like and GC Tfh cell accumulation and maintenance

The second predominant CD4^+^ effector T cell subset that emerges after *Plasmodium* infection are Tfh cells, characterized by the expression of CXCR5, PD-1, and Bcl6 [15]. Tfh cells expand in the spleen in response to *Plasmodium* infection and are critical for producing isotype-switched Abs. Two distinct populations of Tfh cells emerge during infection: Tfh-like (CXCR5^+^PD-1^+^) and GC Tfh (CXCR5^+^PD-1^++^) cells [2]. While the loss of ICOS does not impair the emergence of Tfh-like cells during the early stages of infection with *P. yoelii* (Fig. 2A), their abundance and total numbers are significantly lower than those found in WT mice at day 7 p.i. (Fig. 2A,B). Furthermore, the progression of Tfh-like cells to a GC Tfh cell phenotype was significantly impaired in the *Icos*^-/-^ mice (Fig. 2A, C). Although their numbers were lower, the *Icos^-/-^* Tfh cells expressed Bcl6, a transcription factor that dictates Tfh cell differentiation [16]. Moreover, the MFI for Bcl6 was comparable between WT and *Icos^-/-^* Tfh cells at day 7 p.i. (Fig. 2D).

**FIG. 2.**
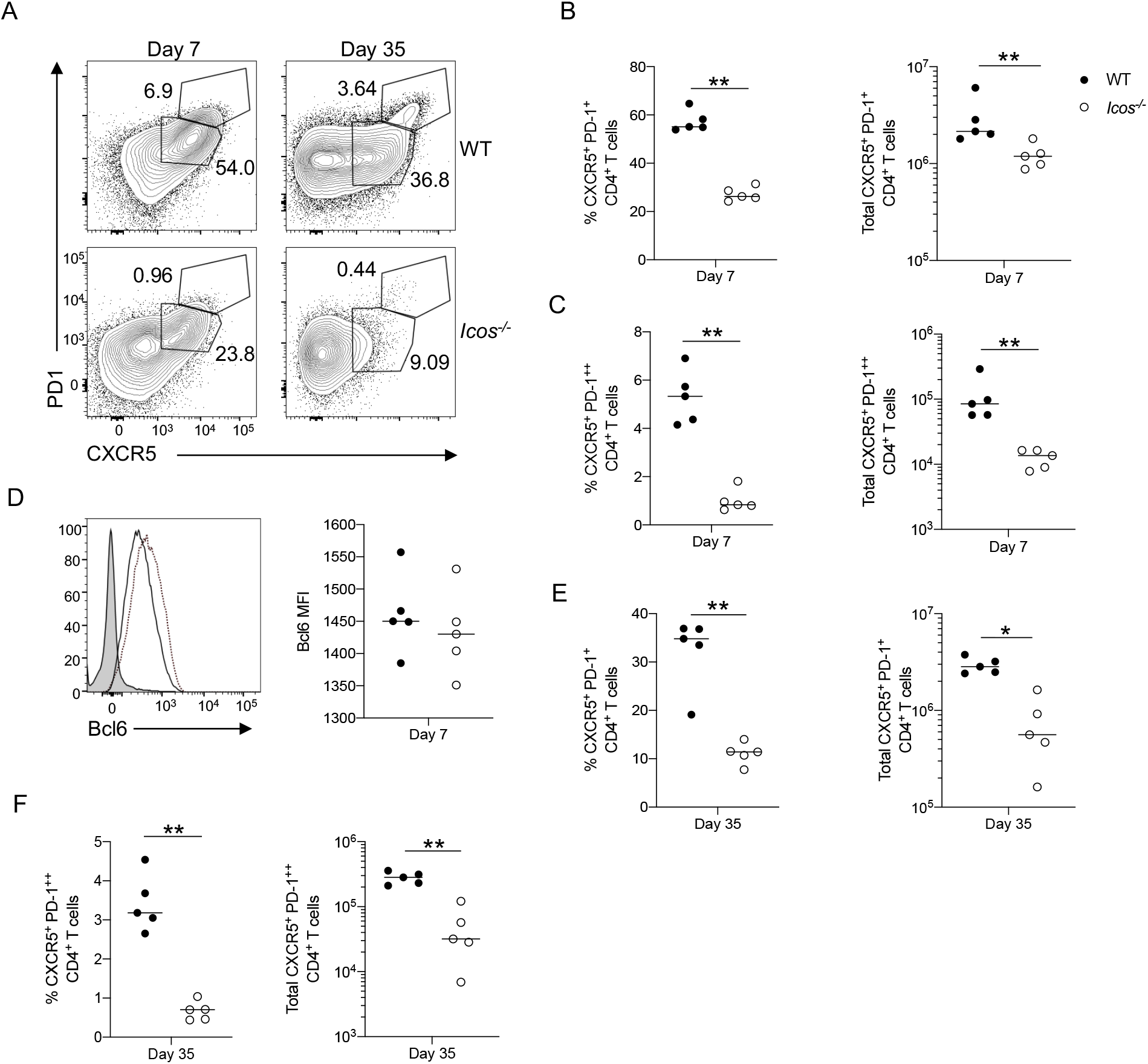
*Icos^-/-^* mice exhibit defects in Tfh-like and GC Tfh cell accumulation and maintenance. WT and *Icos^-/-^* mice were infected i.p. with 10^5^ *P. yoelii* pRBCs. (**A**) Representative flow plots of live activated Tfh-like (CXCR5^+^PD-1^+^) and GC Tfh (CXCR5^+^PD-1^++^) CD4^+^ T cells at day 7 and 35 p.i. from WT and *Icos^-/-^* mice. Frequency and the total number of live activated (**B**) Tfh-like and (**C**) GC Tfh cells at day 7 p.i. (**D**) Histogram and MFI (median) of Bcl6 expression in live CD44^hi^CD11a^+^ CD4^+^ T cells at day 7 p.i. FMO is indicated by the shaded peak, WT by a solid line, and *Icos^-/-^* by a dotted line. Frequency and the total number of live activated (**E**) Tfh-like and (**F**) GC Tfh cells at day 35 p.i. Data are representative of 3-4 independent experiments with five mice per group. A nonparametric Mann-Whitney *t*-test determined significance. \**p* < 0.05, \*\**p* <0.01.

By day 35 p.i., the Tfh-like population in *Icos^-/-^* mice remained significantly diminished compared to WT mice (Fig 2A,E). Also, a distinct GC Tfh cell population was observed in WT mice at day 35, but it was largely absent in *Icos^-/-^* mice (Fig. 2A,F), indicating a failure of *Icos^-/-^* mice to develop and retain this population after resolution of the infection. These results suggest that although Tfh cell accumulation and maintenance are negatively impacted in the absence of ICOS, this receptor is not required for initial Tfh cell formation during *P. yoelii* infection.

### Early Ag-specific Ab production is diminished in the absence of ICOS, but GCs form

Since the Tfh cell response is directly tied to the B cell response, including Ag-specific Ab production, the diminished Tfh cell response in *Icos^-/-^* mice suggests an impairment in the humoral immune response to *P. yoelii* infection in the absence of ICOS. Indeed, early Ag-specific IgM and IgG production was reduced in *Icos^-/-^* mice (Fig. 3A). This observation correlated with a defect in the early plasmablast response (Supplemental Fig. 2A, Fig. 3B). Furthermore, while the frequency of B cells with a GC phenotype (Fas^+^GL-7^+^) was reduced in the absence of ICOS at days 7 and 11, no impact on cell numbers was observed at this time (Supplemental Fig. 2B, C; Fig. 3C,D). On day 11 p.i. the inflammation associated with *P. yoelii* infection results in splenomegaly, causing the white pulp regions to be spread out more from one another and disorganization of the T cell zones (Supplemental Fig. 2D).

**FIG. 3.**
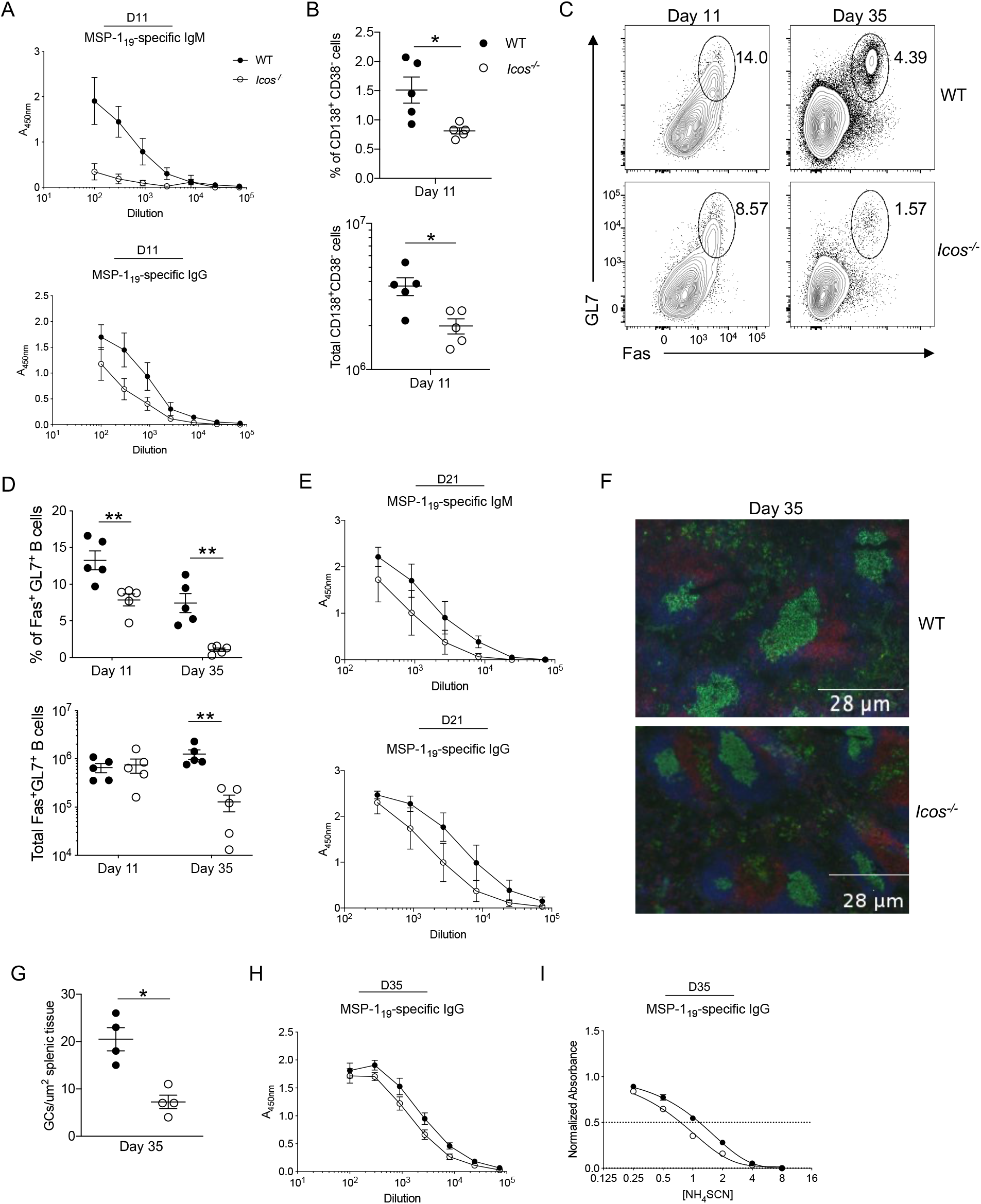
GCs form, and Ag-specific Ab is produced in the absence of ICOS. (**A**) Serum MSP-119–specific IgM and IgG titers were determined by ELISA on day 11 p.i. (**B**) Frequency and the total number of live CD138^+^CD38^-^ plasmablasts at day 11 p.i. (**C**) Representative flow plots of GL7 and Fas expression on live CD19^+^B220^+^ B cells from WT and *Icos^-/-^* mice at days 11 and 35 p.i. (**D**) Frequency and the total number of live GC B cells at the specified time points. (**E**) Serum MSP-119–specific IgM and IgG were determined by ELISA at day 21 p.i. (**F**) 4 μm spleen sections from WT and *Icos^-/-^* mice were stained with CD3*ε* (red), IgD (blue), and GL7 (green) at day 35 p.i. (**G**) Enumeration of GL7^+^ GCs per μm^2^ of splenic tissue at day 35 p.i. (**H**) Serum MSP-119–specific IgG was determined by ELISA at day 35 p.i. (**I**) Normalized absorbance of MSP-119–specific IgG at day 35 p.i. as determined by ELISA. The x-axis (log2) displays increasing molar concentrations of NH4SCN. A nonlinear regression curve fit is shown; the intersection of the dotted line at the normalized absorbance of 0.5 with the curves represents the concentration of NH4SCN needed to elute 50% of the bound IgG from the Ag. Data are representative of 2-3 independent experiments with 3-5 mice per group. (A-D and F) A nonparametric Mann-Whitney *t*-test determined significance. \**p* < 0.05, \*\**p* <0.01. (E, H, and I) An unpaired, two-tailed Student *t* test was used to determine significance.

Nevertheless, B cell follicles are present, including some that stain positive for GL7, denoting the initial formation of GC structures within the spleen at this time. These early GC structures were apparent in the *Icos^-/-^* mice. However, most of the GL7 staining occurred outside the follicle in the spleen of WT and *Icos^-/-^* mice, depicting the extrafollicular response, which dominates at this time during the infection. The difference in Ab production was short-lived, as by day 21 p.i. the gap in Ag-specific IgM and IgG production decreased as titers rose in both groups of mice (Fig. 3E).

With the resolution of the infection, the frequency of GC B cells decreased on average in the spleen of WT and *Icos^-/-^* mice by day 35, with a significantly higher frequency found in WT mice (Fig. 3C, D). However, unlike day 11, substantially fewer GC B cells were found in the spleen of *Icos^-/-^* mice at day 35 (Fig. 3D), suggesting a defective GC response. However, GC structures were prevalent in the spleen of WT and *Icos^-/-^* mice at day 35 p.i. (Fig. 3F). This was surprising given the significant reduction in GC Tfh and B cell numbers in *Icos^-/-^* mice at this time. However, enumeration of the GCs indicated that significantly fewer of these structures were present in *Icos^-/-^* mice (Fig. 3G).

Similar to day 21, the difference in serum Ag-specific IgG production was minimal between WT and *Icos^-/-^* mice (Fig. 3H and Supplemental Fig. 3A). As the GC response progresses, the affinity of Ab for binding Ag should increase. Therefore, the affinity maturation of Ag-specific Abs from WT and *Icos^-/-^* mice was compared to understand whether the GCs are functional in the *Icos^-/-^* mice. The affinity of the IgG Ab specific for MSP-1 and AMA-1 was not significantly different between WT and *Icos^-/-^* mice, though it was lower for MSP-1 in the *Icos^-/-^* mice (Fig. 3I and Supplemental Fig. 3B). Together, these results suggest that an early defect in Ag-specific Ab production likely contributed to the delayed resolution of the infection in *Icos^-/-^* mice, which may in part be due to the reduction in Tfh cell accumulation. However, while Tfh cells were significantly reduced in the absence of ICOS, the data indicate that ICOS is dispensable for GC formation in response to *P. yoelii* infection. Thus, the ability of *Icos^-/-^* mice to form these GC structures, albeit at a lower overall quantity, and continue their production of Ag-specific Abs at titers seen in WT mice likely allowed them to control and clear the infection eventually.

### Blocking ICOS:ICOSL interactions during infection do not impair GC formation

To determine if ICOS:ICOSL interactions during *P. yoelii* infection were specifically responsible for the phenotype observed in the *Icos^-/-^* mice, WT mice were treated with an anti-ICOSL monoclonal Ab or an isotype control Ab starting one day before infection and every three days after. Similar to the *Icos^-/-^* mice and previous findings with ICOSL blockade, resolution of the infection with *P. yoelii* was delayed following anti-ICOSL treatment (Fig. 4A). The delay in parasite clearance correlated with a defect in the appearance of GC Tfh cells but not Tfh-like cells in the anti-ICOSL treated mice (Fig. 4B). While the frequency of Tfh-like cells was similar between groups, the overall number of Tfh-like cells was reduced in WT mice in response to ICOSL blockade and comparable to numbers observed in *Icos*^-/-^ mice (Fig. 4C). Likewise, the overall number of GC Tfh cells was reduced in the anti-ICOSL treated mice compared to the isotype control treated mice (Fig. 4D). As expected, the reduction in Tfh cell accumulation with ICOSL blockade negatively impacted the GC B cell response. The frequency and number of Fas^+^GL-7^+^ GC B cells were decreased compared to WT mice that received the isotype control Ab. However, the decline in GC B cells was not as significant as that observed in the *Icos^-/-^* mice (Fig. 4E,F). Despite the defect in Tfh cell and GC B cell accumulation, GC structures were apparent in the spleen of the anti-ICOSL treated mice, just as they were evident in the spleen of *Icos^-/-^* mice (Fig. 4G). However, the number of visible GC structures was reduced with anti-ICOSL treatment (Fig. 4H). Yet, no difference in MSP-1–specific IgG production or affinity for binding Ag was detected between the anti-ICOSL and isotype treated mice in response to *P. yoelii* infection (Fig. 4I). Hence, the results seen with the anti-ICOSL blockade in WT mice resembled those observed in *Icos^-/-^* mice after *P. yoelii* infection and indicated that ICOS signaling, while important for producing Tfh cells, is not required for the generation of a humoral response capable of resolving a *P. yoelii* infection.

**FIG. 4.**
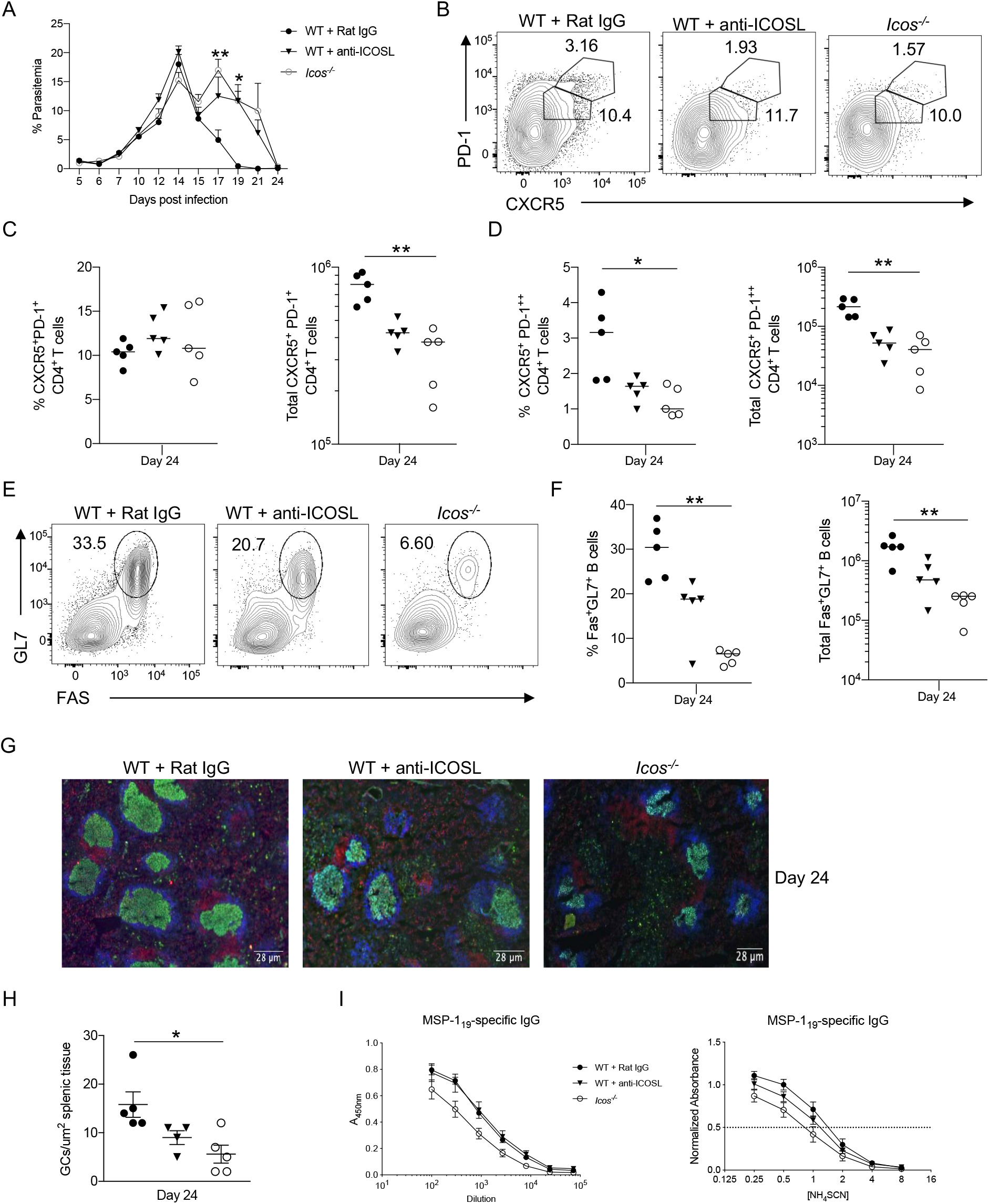
GC formation is not impaired with ICOSL blockade. WT mice were injected i.p. with 200 μg of anti-ICOSL or rat IgG isotype control Ab starting one day prior to infection and every three days afterward until the time of sacrifice. (**A**) Representative parasitemia curve as determined by flow cytometry. (**B**) Representative flow plots of live activated Tfh-like and GC Tfh cells at day 24 p.i. Frequency and total number of live activated (**C**) Tfh-like and (**D**) GC Tfh cells at day 24 p.i. (**E**) Representative flow plots of GL7 and Fas expression on live CD19^+^B220^+^ B cells at day 24 p.i. (**F**) Frequency and the total number of Fas^+^GL7^+^ GC B cells at day 24 p.i. (**G**) 4 μm spleen sections from WT and *Icos^-/-^* mice were stained with CD3*ε* (red), IgD (blue), and GL7 (green) at day 24 p.i. (**H**) (I) Serum MSP-119–specific IgG was determined by ELISA on day 24 p.i. (left). Normalized absorbance of MSP-119–specific IgG at day 24 p.i. as determined by ELISA displayed as a nonlinear regression curve fit (right). The x-axis (log2) displays increasing molar concentrations of NH4SCN. Data are representative of two independent experiments with 3-5 mice per group. A two-way ANOVA determined significance with a post hoc Holm-Sidak’s multiple comparison test. \**p* < 0.05, \*\**p* < 0.01.

### ICOS is necessary for optimal expression of molecules that support GC Tfh cell function and interactions with B cells

To determine if the GC Tfh cells present in *Icos^-/-^* mice can support the GCs that form, they were examined for the expression of key genes that promote and sustain the GC response. Sort-purified GC Tfh cells (Supplemental Fig. 4A) recovered from *Icos^-/-^* mice 11 days after infection with *P. yoelii* showed a reduction in *il4*, *il21*, *ifng*, *tbx21*, *bcl6,* and *cd40l* expression compared to GC Tfh cells derived from WT mice (Fig. 5A). A similar trend in the expression of these genes was noted in Tfh-like cells sorted from *Icos^-/-^* mice compared to those recovered from WT mice, except for *il4*, which was not reliably detected in the Tfh-like cells (Fig. 5B).

**FIG. 5.**
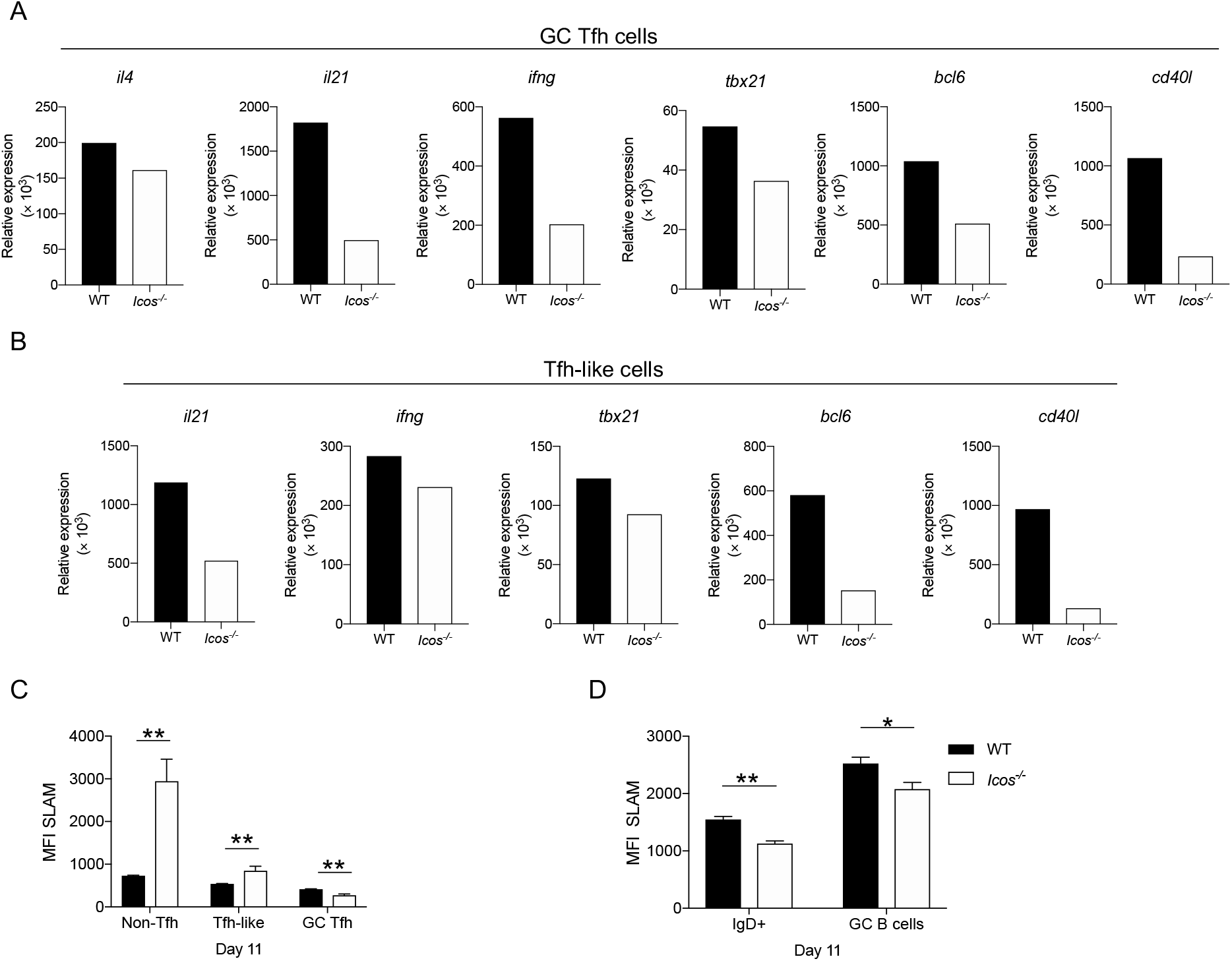
GCs have functionality defects in the absence of ICOS. Relative expression of *il4*, *il21*, *ifng*, *tbx21*, *bcl6*, and *cd40l* in **(A)** GC Tfh cells and (B) Tfh-like cells (excluding *il4*) at day 11 p.i. as determined by real-time quantitative PCR. Data were normalized to *hprt*, and the 2^-DCt^ method was used to calculate fold changes. **(C)** MFI of SLAM expression on live activated non-Tfh (CXCR5^-^PD-1^+/-^), Tfh-like, and GC Tfh CD4^+^ T cells at day 11 p.i. (D) MFI of SLAM expression on live CD19^+^B220^+^ IgD^+^ or Fas^+^GL7^+^ B cells at day 11 p.i. Data are representative of two independent experiments with five mice per group. A nonparametric Mann-Whitney *t-*test determined significance. \**p* < 0.05, \*\**p* <0.01.

SLAM (CD150), the prototypic member of the SLAM family of receptors that is expressed by activated T cells, is involved in interactions between GC Tfh and B cells that promote adhesion and bidirectional signaling. Here we note that its expression is significantly elevated on non-Tfh and Tfh-like cells in *Icos^-/-^* mice at day 11 after *P. yoelii* infection (Fig. 5C). However, amongst the GC Tfh cells, SLAM expression is appreciably reduced in the absence of ICOS. Furthermore, while SLAM is upregulated on GC B cells from WT and *Icos^-/-^* mice compared to IgD^+^ B cells, its expression is significantly reduced on both populations of B cells in *Icos^-/-^* mice (Fig. 5D). Moreover, the reduced expression of SLAM on GC Tfh and B cells remained consistent at day 36 p.i. (Supplemental Fig. 3B,C). Hence, interactions between GC Tfh and B cells may be compromised within the GC of *Icos^-/-^* mice.

Light zone B cells are known to upregulate MHC class II, CD86, and CD83 expression to facilitate their interaction with CD4^+^ T cells in the GC [17]. However, it is unclear if the loss of ICOS modulates their expression. Therefore, B cells were examined for changes in the expression of these proteins during and after recovery from *P. yoelii* infection. Of these proteins, only CD86 expression was significantly reduced on GC B cells without ICOS, which was only at day 36 p.i. (Supplemental Fig. 4D,E). Alternatively, IgD^+^ B cells displayed reduced amounts of MHC class II and CD86 in the absence of ICOS at days 11 and 36, suggesting that interactions between T and B cells outside the GC may also be compromised in the absence of ICOS. Overall, the loss of ICOS induced phenotypic changes in GC Tfh and GC B cells, suggesting that functional defects exist in those GCs present in *Icos^-/-^* mice.

### GCs deteriorate faster in the absence of ICOS

Given the potential functional defects in the GC response in *Icos^-/-^* mice, we hypothesized that the GC structures present in the spleen of these mice at day 35 would dissociate sooner than those seen in WT mice. GC Tfh and B cells are still abundantly present in the spleen of WT mice at day 70 p.i. However, these populations remained significantly deficient in the *Icos^-/-^* mice at day 70 (Fig. 6A, B). Furthermore, by day 70, GC structures were no longer observed in the spleen of *Icos^-/-^* mice, but they were still present in WT mice, although smaller in size than day 35 (Fig. 6C, D). Surprisingly, MSP-1– and AMA-1–specific IgG production did not vary greatly between WT and *Icos^-/-^* mice (Fig. 6E, Supplemental Fig. 3C). As GC B cells undergo continual rounds of proliferation and somatic hypermutation within the GC, the affinity of the BCR on the daughter B cells for binding Ag and the subsequent Ab produced by plasma cells that emerge from the GC should increase. Indeed, the affinity of the MSP-1– and AMA-1–specific IgG derived from WT mice for binding its cognate Ag increased from day 35 to day 70 (Fig. 3I, Fig. 6E and Supplemental Fig. 3B, D). However, no improvement in the affinity of the Ag-specific IgG for binding these two Ags was observed in the absence of ICOS signaling. This result points to a defect in somatic hypermutation in the GC of *Icos^-/-^* mice. Overall, the data suggested that although ICOS expression was not necessary to promote GC formation, it was important for preserving the GC function and integrity over time after clearance of the infection.

**FIG. 6.**
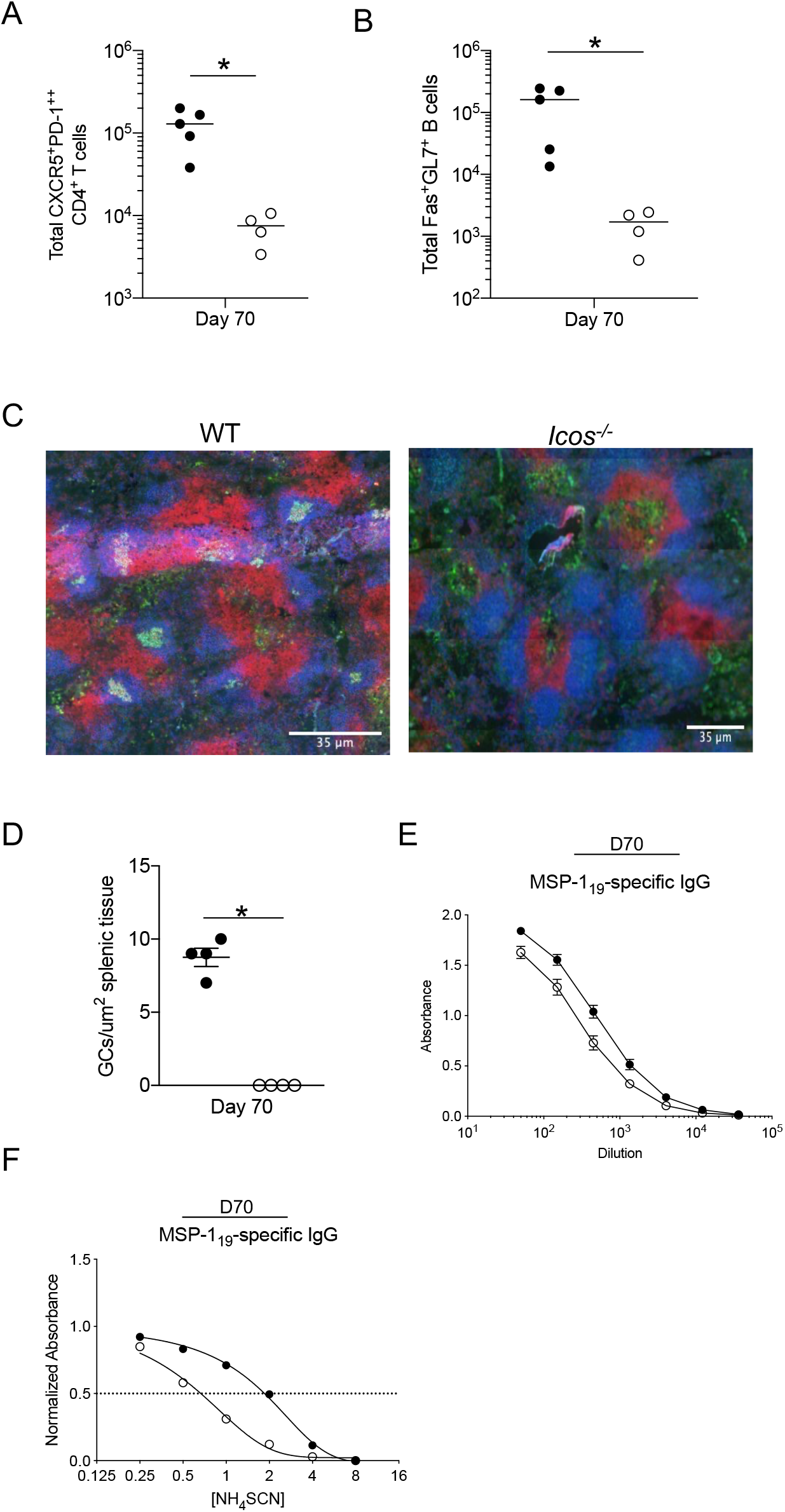
GCs deteriorate faster in the absence of ICOS. Total number of (**A**) live activated CXCR5^+^PD-1^++^ GC Tfh cells and **(B)** live CD19^+^B220^+^ Fas^+^ GL7^+^ GC B cells at day 70 p.i. **(C)** 4 μm spleen sections from WT and *Icos^-/-^* mice were stained with CD3*ε* (red), IgD (blue), and GL7 (green) at day 70 p.i. **(D)** Enumeration of GC structures per μm^2^ of splenic tissue at day 70 p.i. **(E)** Serum MSP-119–specific IgG was determined by ELISA at day 70 p.i. **(F)** Normalized absorbance of MSP-119–specific IgG at day 70 p.i. as determined by ELISA displayed as a nonlinear regression curve fit. The x-axis (log2) displays increasing molar concentrations of NH4SCN. Data are representative of two independent experiments with 4-5 mice per group. (A-D) A nonparametric Mann-Whitney *t*-test determined significance. \**p* < 0.05. (E-F) An unpaired, two-tailed Student *t* test was used to determine significance.

### Reduction in Ag load does not impede GC formation in *Icos^-/-^* mice

Given the importance of ICOS in Tfh cell differentiation and the formation of GCs, our findings that these processes still occur at a reduced capacity in the absence of ICOS during *P. yoelii* infection raises the question as to what instead drives this response. One candidate is excess Ag, as the increased availability of Ag can lead to Tfh cell formation in the absence of ICOS [10, 11]. Infection of mice with *P. yoelii* results in high parasite loads in the blood (>20% of RBCs infected). Therefore, one potential explanation for the ability of *Icos^-/-^* mice to form Tfh cells and GCs is that the high Ag load associated with this infection allows the immune system to bypass the requirement for ICOS in Tfh cell induction and GC formation. To test this idea, WT and *Icos^-/-^* mice were drug-cured using the anti-malarial drug atovaquone (AV) to reduce the Ag load associated with *P. yoelii* infection (Fig. 7A). AV treatment prevented parasitemia from reaching 5% in WT and *Icos^-/-^* mice, and the infection was cleared by day 14 p.i. Meanwhile, peak parasitemia exceeded 20% in the non-treated groups, with the *Icos^-/-^*mice again showing a defect in their ability to control the infection (Fig. 7B).

**FIG. 7.**
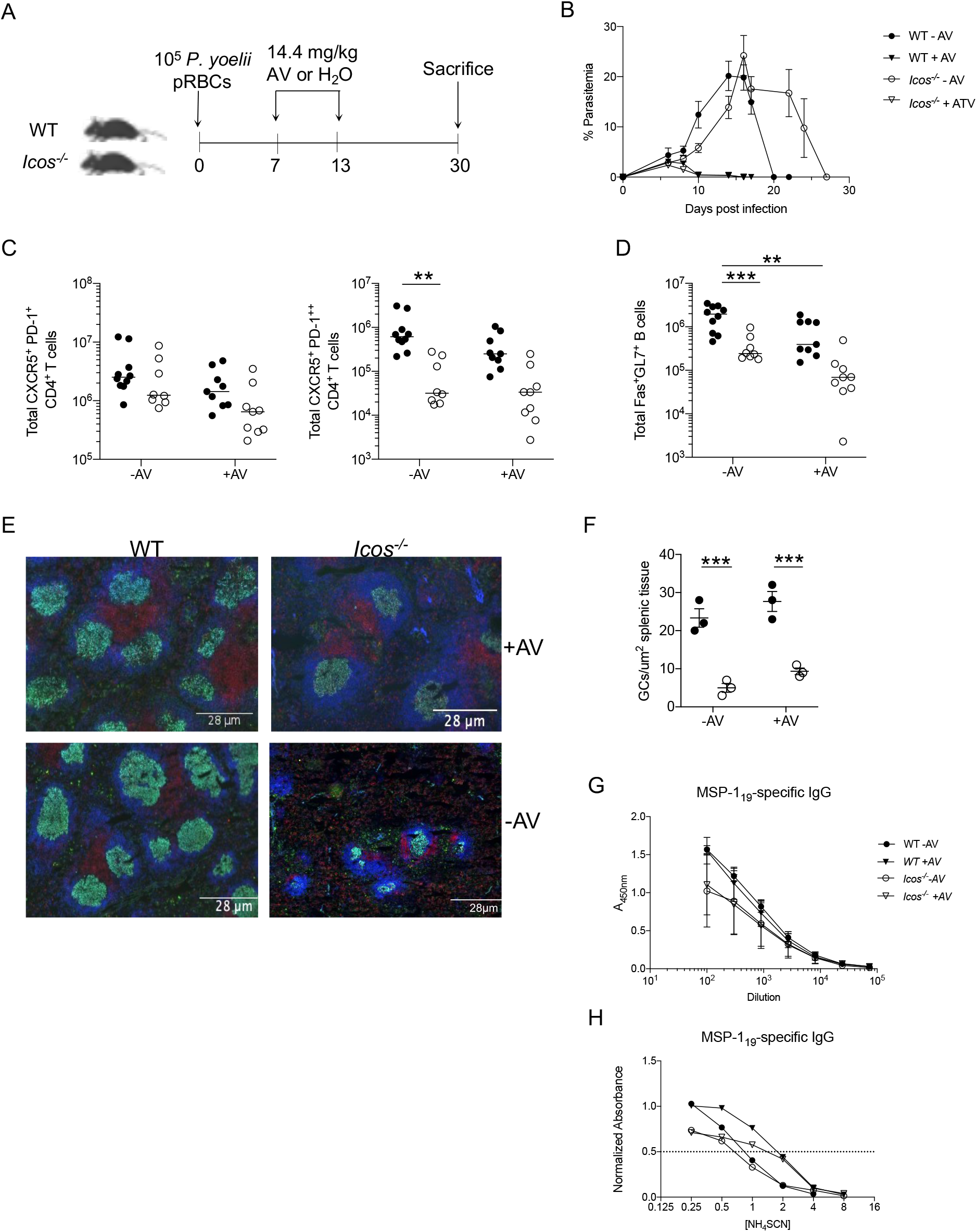
Reduction in Ag load does not impede GC formation in *Icos^-/-^* mice. **(A)** Experimental model. WT and *Icos^-/-^* mice were infected i.p. with 10^5^ *P. yoelii* pRBCs. From days 7 to 13 p.i. mice were treated i.p. with 14.4 mg/kg of atovaquone or sterile water. Mice were sacrificed on day 30 p.i. **(B)** Representative parasitemia curve as determined by flow cytometry. **(C)** Total numbers of live activated Tfh-like (CXCR5^+^PD-1^+^) and GC Tfh (CXCR5^+^PD-1^++^) cells at day 30 p.i. **(D)** Total numbers of live CD19^+^B220^+^ Fas^+^GL7^+^ B cells at day 30 p.i. **(E)** 4 μm spleen sections from WT and *Icos^-/-^* mice were stained with CD3*ε* (red), IgD (blue), and GL7 (green) at day 30 p.i. **(F)** The total number of GCs per μm^2^ of splenic tissue at day 30 p.i. **(G)** Serum MSP-119–specific IgG was determined by ELISA at day 30 p.i. **(H)** Normalized absorbance of MSP-119–specific IgG at day 30 p.i. as determined by ELISA displayed as a nonlinear regression curve fit. The x-axis (log2) displays increasing molar concentrations of NH4SCN. Data are pooled from two independent experiments. A two-way ANOVA determined significance with a post hoc Holm-Sidak’s multiple comparison test. \*\**p* < 0.01, \*\*\**p* < 0.001.

Assessment of the accumulation of Tfh and GC B cell populations at day 30 p.i. indicated that regardless of treatment with AV, GC Tfh and B cell accumulation was impaired in the absence of ICOS compared to WT mice (Fig. 7C, D). Although, the difference in cell numbers for each population was reduced with AV treatment, primarily due to a greater effect of drug treatment on WT cell numbers. Based on these results, we anticipated finding GC structures in the spleen of the AV treated *Icos^-/-^* mice, similar to the non-treated controls. Indeed, this was the case, as GCs were found throughout the spleen of *Icos^-/-^* mice regardless of whether they received AV (Fig. 7E). Furthermore, significantly fewer GCs were present in the spleen of all *Icos^-/-^* mice, with similar numbers seen between those that did and did not receive AV (Fig. 7F). Thus, reducing the Ag load in *Icos^-/-^* mice did not further impact Tfh cell or GC B cell formation after *P. yoelii* infection.

Furthermore, MSP-1–specific IgG production was comparable between WT or *Icos^-/-^* mice that did or did not receive AV, with titers trending lower in the *Icos^-/-^* groups (Fig. 7G). While no difference in Ab affinity was seen between WT and *Icos^-/-^* mice treated with or without AV, the affinity of IgG for binding MSP-1 was lower in the non-treated mice compared to those that received AV (Fig. 7H). The reduced inflammation seen in the spleen of AV treated mice likely allowed GC reactions to proceed at a faster rate than in the non-treated mice, as evident by the more prominent and organized GCs and T cell zones seen in the spleen of the non-treated mice at day 21 (Supplemental Fig. 5). Thus, although Ag load was reduced with AV treatment, sufficient Ag was available to promote Tfh accumulation, GC formation, and drive Ab production even in the absence of ICOS.

### Secondary GCs do not form in the absence of ICOS

Given the ability of the *Icos^-/-^* mice to resolve their primary infection with *P. yoelii* and form temporary GC structures, we were interested in determining if the humoral response generated during the primary infection was sufficient to afford protection upon re-challenge. Re- infection of C57BL/6 mice 70-95 days after primary infection with *P. yoelii* results in a blood parasitemia that is below the limit of detection on a Giemsa-stained thin blood smear, unlike re- infection with *P. chabaudi*, which causes a persistent infection but of lower magnitude than the primary infection [18], indicating that WT mice are afforded protection against secondary infection with *P. yoelii*. However, the immune response is activated after re-challenge with *P. yoelii* based on increases in Ab titers and an expansion in Ab-secreting cells (Figure 8A,B). However, like what we observed in the *P. chabaudi* model after secondary infection [19] the secondary humoral response was impaired without ICOS. Although, no parasites were ever detectable in the blood smears, indicating that the presence of Ag-specific Abs, though of a lower affinity, was sufficient to protect these mice.

**FIG. 8.**
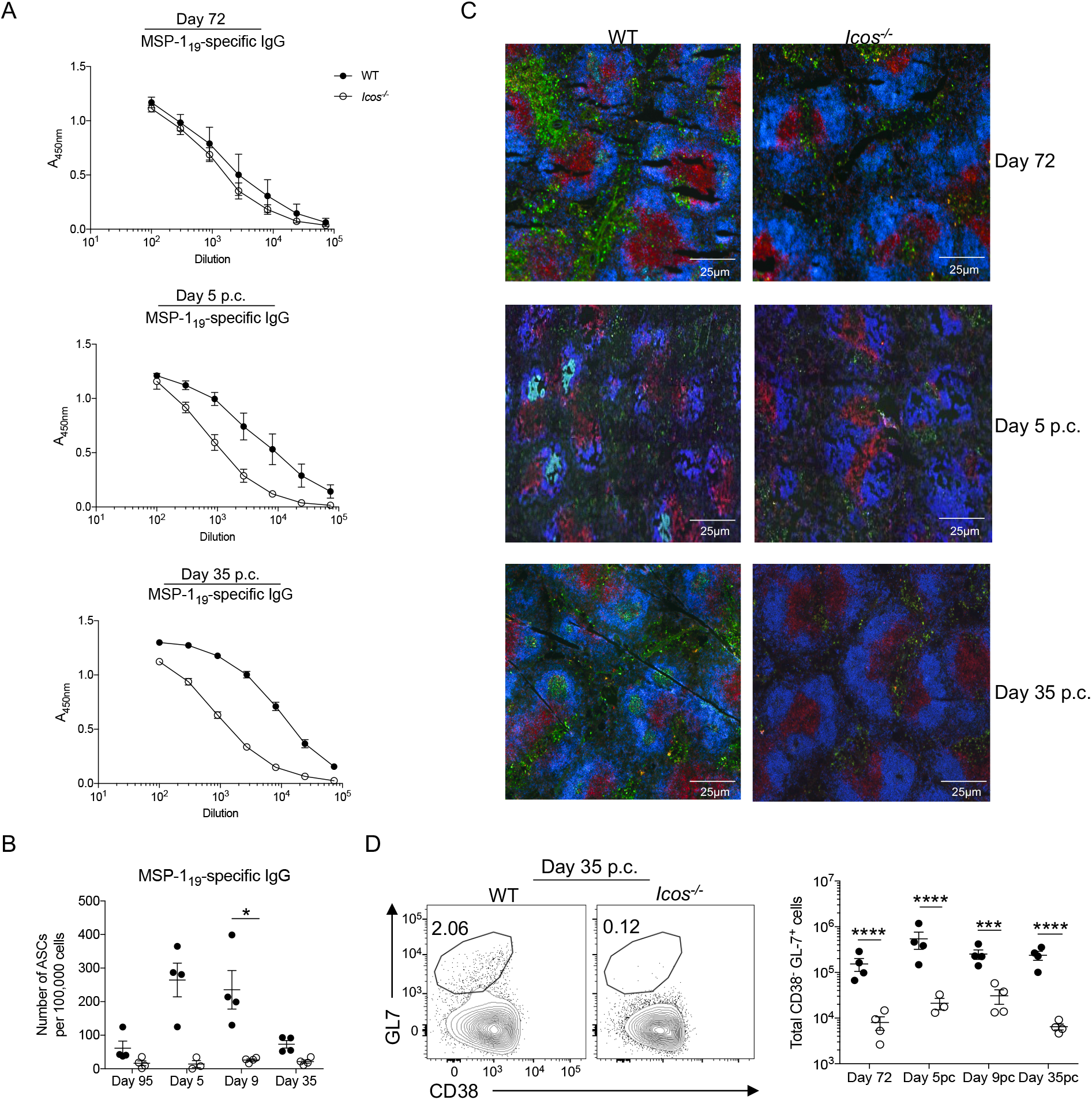
Secondary GCs do not form in the absence of ICOS. (**A**) Measurement of MSP-119–specific IgG in the serum of WT and *Icos^-/-^* mice at days 72 p.i., 5 p.c., and 35 p.c. as determined by ELISA. (**B**) The number of IgG^+^ ASCs per 10^5^ splenocytes specific for MSP-119 as determined by ELISpot at the indicated time points. **(C)** 4 μm spleen sections from WT and *Icos^-/-^* mice were stained with CD3*ε* (red), IgD (blue), and GL7 (green) at days 72 p.i., 5 post challenge (p.c.), and 35 p.c. **(D)** Representative plots of CD38 and GL-7 expression on live CD19^+^B220^+^ B cells from WT and *Icos^-/-^* mice at day 35 p.c. Total numbers of CD38^-^GL7^+^ B cells at indicated time points. Data are representative of two independent experiments with 3-5 mice per group. (A) An unpaired, two-tailed Student *t* test was used to determine significance. (B and D) A nonparametric Mann-Whitney *t-*test determined significance. \**p* < 0.05, \*\*\**p* < 0.001, \*\*\*\**p* < 0.0001.

To determine the capacity of *Icos^-/-^* mice to form secondary GCs after re-infection, spleen sections from WT and *Icos^-/-^* mice were examined 5 and 35 days after re-infection. Residual primary GCs are still present in the spleen of WT but not *Icos^-/-^* mice prior to re-challenge (Fig. 8C). Unlike secondary infection with *P. chabaudi*, which induces an inflammatory response capable of dissociating GC structures within the spleen, GC structures are visible in the spleen of WT mice 5 days after re-infection with *P. yoelii* (Fig. 8C). However, no GC structures were observed in the spleen of *Icos^-/-^* mice 5 days after re-challenge. Moreover, while GCs are visible and abundant in the spleen of WT mice at day 35 p.c., de novo formation of GCs is not seen in the spleen of *Icos^-/-^* mice. Indeed, flow cytometry analysis confirmed that GC B cells were present in the spleen of WT mice, expanding after re-challenge. At the same time, they were significantly lower in *Icos^-/-^* mice before and after re-challenge, although their cell numbers expanded at day 5 p.c. (Fig. 8D). Together these data suggest that ICOS plays an important role in supporting secondary humoral responses. Other factors cannot compensate for the loss of ICOS in these processes, particularly the de novo formation of new GCs after re-infection.

## Discussion

The data shown here demonstrate that the co-stimulatory molecule ICOS is not necessary for CD4^+^ T cells to express markers associated with a Tfh cell phenotype, nor is it required to form GC structures in response to infection with *P. yoelii*. The ability of *Icos^-/-^* mice to form GCs here is opposite of our findings in a *P. chabaudi* model [5], indicating there are differences in the signals needed to generate their structures in each model. However, ICOS was crucial for maintaining these structures over time and CD4^+^ T cells with a GC Tfh cell phenotype, the latter finding resembling the *P. chabaudi* model [5]. *Icos^-/-^* mice could produce ASCs and parasite-specific Abs, albeit at lower quantities early during the infection, which impacted the ability of the mice to control and resolve their infection. Whether these ASCs were directly derived from the GC, a product of the extrafollicular response, or a combination of each pathway remains unclear. Though the lack of affinity maturation in the parasite-specific Abs suggests a functional defect in the GC, it does not rule out that the GCs are incapable of supporting long-lived plasma cell formation before their dissociation in the absence of ICOS signaling. However, it is less likely that these GCs are a source of long-lived ASCs in these mice.

After observing the formation of GCs in *Icos^-/-^* mice, we initially hypothesized that the high Ag load that accompanies a *P. yoelii* infection afforded these mice the ability to form GCs in the absence of ICOS. High Ag load in other cases can promote GC reactions. For instance, a direct correlation was observed between the GC response and Ag load where increasing the amount of Ag delivered resulted in an increase in the Tfh and GC B cell response in an OVA immunization model [20]. In the same report, repeated injection of OVA to provide a constant supply of Ag also led to an increase in the frequency of Tfh and GC B cells compared to mice that received a single injection with OVA. Two related studies demonstrated that chronic viral infection with LCMV, and hence continued presence of Ag, yielded GC responses of higher cellularity than acute infection with LCMV. Chronic infection led to an increase in memory B cell and ASC output, efficient selection of hypermutated clones, and higher titers of neutralizing Abs [21, 22], indicating that the humoral response excels when continuously faced with high amounts of viral Ag. However, even though AV significantly reduced the parasite burden, enough Ag was available to promote a humoral response and the formation of GC structures in WT and *Icos^-/-^* mice. Thus, a high Ag load is not required to enable a GC response with *P. yoelii* infection.

Ag load is not the only dictator of Ab development during infection, as Chan and associates demonstrated that during experimental *P. falciparum* infection in humans Ab development was not reliant on Ag load [23], which contrasts with other findings [24, 25, 26]. Rather there was a correlation between Ab development and Th2-cTfh cell populations. [23]. Moreover, the avidity of the TCR for binding peptide:MHC complexes was also suggested to provide early instructions for promoting Tfh cell fate decisions. However, the findings in various models are inconsistent with one another, with some studies suggesting that strong TCR avidity for peptide:MHC promotes Tfh cell development [27, 28, 29]. In contrast, others concluded the opposite [30, 31] or found no effect on Tfh cell development [32, 33]. A more recent finding suggested that basal, tonic TCR signaling could contribute to these diverse results and indicated that CD4^+^ T cells that receive low tonic TCR signaling favor Tfh cell differentiation. In contrast, strong tonic signaling inhibits it [34].

Other factors that influence GC reactions during infection that could account for the loss of ICOS signaling include the co-stimulatory receptor OX40. OX40 is expressed on activated T cells and provides pro-survival signals that promote humoral immunity [35]. Stimulation of OX40 with an Ab agonist during vaccination in a murine model resulted in an enhanced expansion of CD4^+^ effector T cells and Ag-specific IgG [36]. Also, the ligation of OX40 leads to the generation of memory Th1 and Tfh CD4^+^ T cells [14]. Thus, this co-stimulatory molecule might act in an ICOS deficient environment during *P. yoelii* infection to still allow for the generation of Tfh cells, albeit their generation is significantly decreased to what is observed in WT mice. Determining if OX40 or possibly another co-stimulatory molecule is offsetting the absence of ICOS signaling during *P. yoelii* infection requires further study.

The intracellular adaptor protein SAP (SLAM associated protein) is another factor that could promote GC reactions in the absence of ICOS. This protein interacts with the cytoplasmic tails of the SLAM family of receptors and mediates downstream signaling [37]. SAP is known to modulate interactions between Tfh and B cells, which lead to the promotion of GCs, and its intrinsic expression specifically in CD4^+^ T cells is critical for GC formation [38]. Like ICOS, SAP is required to maintain Tfh cells and promote their localization to the GC [39]. However, the initial accumulation of Tfh cells is not impeded by the absence of SAP in response to *P. chabaudi* [40] or LCMV infection [41] or immunization [42], phenotypes that resemble the one observed in *Icos^-/-^* mice infected with *P. chabaudi* [5]. Moreover, SAP deficient mice can form GCs structures in the spleen and mount an Ag-specific IgG response after *P. chabaudi* infection [40], mimicking the phenotype observed here in response to *P. yoelii* infection.

SLAM family receptors are expressed on T and B cells and bind in a homophilic manner. We noted higher expression of SLAM (CD150) on non-Tfh and Tfh-like cells in *Icos^-/-^* mice compared to WT mice. Perhaps, the increased expression of SLAM on these populations of activated T cells aided in accounting for the absence of ICOS signaling, allowing for early Tfh cell development and promoting GC formation. In contrast, we observed a decrease in SLAM expression on GC Tfh and B cells. Further supporting the idea that while additional factors can contribute to the ability of CD4^+^ T cells to initially display markers associated with a Tfh cell fate in the absence of ICOS, eventually, the T cells reach a critical junction in their development where ICOS signaling is required to sustain their Tfh cell fate.

Although GCs formed in *Icos^-/-^* mice during *P. yoelii* infection, these structures were reduced in overall number, and those present deteriorated more quickly in these animals. This suggests that ICOS:ICOSL interactions are necessary for the continued maintenance of these structures in this model. There are other cases in which GCs can initially form but are not maintained long-term due to the loss of important signals such as IL-21 from GC Tfh cells to GC B cells within the GC reaction [43, 44]. Specifically, IL-21 is crucial for GC B cell differentiation and proliferation [45]. Our data show that IL-21 transcript levels are decreased in GC and Tfh-like cells from *Icos^-/-^* mice suggesting that the GC reaction is not maintained during infection. Furthermore, the GCs that form in the *Icos^-/-^* mice may be the product of a thymus-independent response, based on the lack of affinity maturation and the inability of these mice to maintain these structures over time. However, this is unlikely given the lack of GC formation seen in T-cell deficient mice [46] and the absence of GC B cells and class-switched Ab production in CD4-specific Bcl6 knockout mice in response to *Plasmodium* infection [40].

Moreover, upon homologous re-challenge with *P. yoelii*, GCs did not reform in *Icos^-/-^* mice. However, these structures were observed in WT mice, indicating that ICOS signaling is necessary to promote GC formation during secondary infection. This observation is in accordance with what was previously reported in other models [4, 19].

In terms of the differences in GC formation in *Icos^-/-^* mice infected with *P. chabaudi* or *P. yoelii* infection, while a high Ag load was not responsible for GC formation in *Icos^-/-^* mice after *P. yoelii* infection, another explanation could lie with the Th1 response. During *P. chabaudi* infection, *Icos^-/-^* mice had an enhanced CD4^+^ Th1 response, which increased IFN-*γ* production at early timepoints. In contrast, *Icos^-/-^* mice during *P. yoelii* infection had a CD4^+^ Th1 response comparable to WT mice. Thus, it is possible that the enhanced CD4^+^ Th1 response played a role in the constriction of GC formation in the *P. chabaudi* model, thereby preventing the formation of any GC structures in these mice.

Overall, the data presented here indicate that ICOS signaling is unnecessary for *P. yoelii* infection clearance and survival. It is also dispensable for GC formation during primary *P. yoelii* infection. However, ICOS signaling is necessary for the maintenance of GCs after the resolution of the infection, and secondary infection is not sufficient to drive their reformation in the absence of ICOS. The data also indicate that ICOS signaling is not crucial for protection during secondary infection but is important for enhancing the secondary humoral response. Further studies to elucidate how GC structures form in the absence of ICOS could shed light on other critical factors that drive the formation and maintenance of these structures that are necessary for the generation of high affinity Abs. Understanding the intricacies of the GC reaction is critical to our ability to develop an efficacious vaccine that can provide individuals with protection and long-lived immunity against malaria.

## Acknowledgments

*Plasmodium yoelii* strain 17X was obtained through BEI Resources, NIAID, NIH: MRA-749, contributed by D. Walliker. We also thank Dr. James Burns Jr. (Drexel University College of Medicine) for providing the recombinant MSP-142 and AMA-1 proteins. Special thanks to Andrea Harris for her technical assistance as part of the flow cytometry core.

## Author Contribution

Conceived and designed the experiments: JSS, LEL, KAO. Performed the experiments: LEL, KAO, EN, CLF. Analyzed the data: LEL, KAO, JSS. Wrote the paper: JSS, KAO.

---

**Supplemental FIG. 1.**
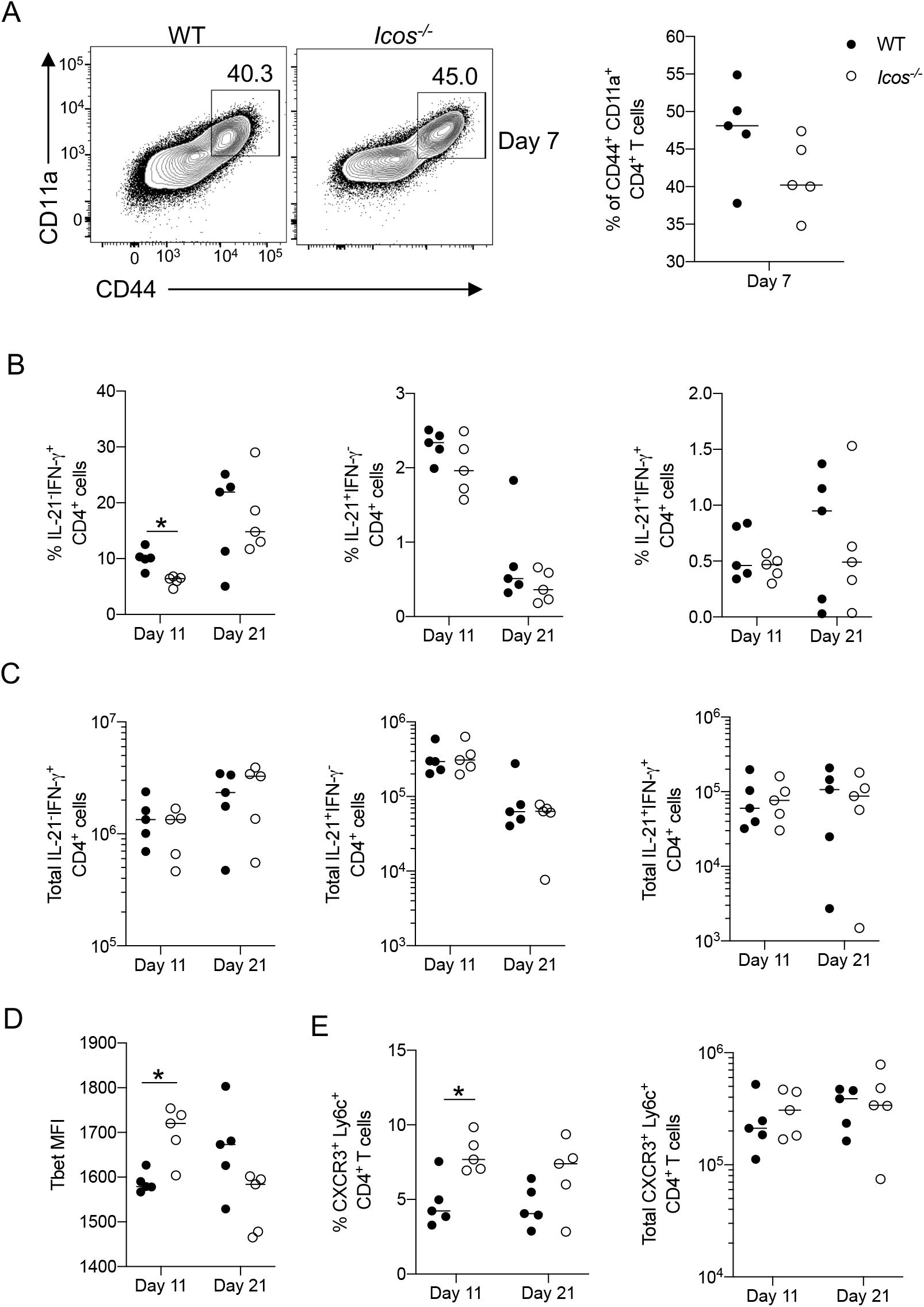
WT and *Icos^-/-^* mice were injected i.p. with 10^5^ *P. yoelii* pRBCs, and splenocytes were analyzed on days 7, 11, and 21 p.i. (**A**) Representative flow plots and frequency of live activated CD4^+^ T cells at day 7 p.i. (**B**) Frequency and (**C**) total number of IL-21^-^IFN-*γ*^+^, IL-21^+^IFN-*γ*^-^, and IL-21^+^IFN-*γ*^+^ CD4^+^ T cells at days 11 and 21 p.i. (**D**) MFI (median) of Tbet expression in live activated CD4^+^ T cells at days 11 and 21 p.i. (**E**) Frequency and the total number of live activated CXCR3^+^ Ly6c^+^ CD4^+^ T cells at days 11 and 21 p.i. Data are representative of 3-4 independent experiments with five mice per group. A nonparametric Mann-Whitney *t*-test determined significance. \**p* < 0.05.

**Supplemental FIG. 2.**
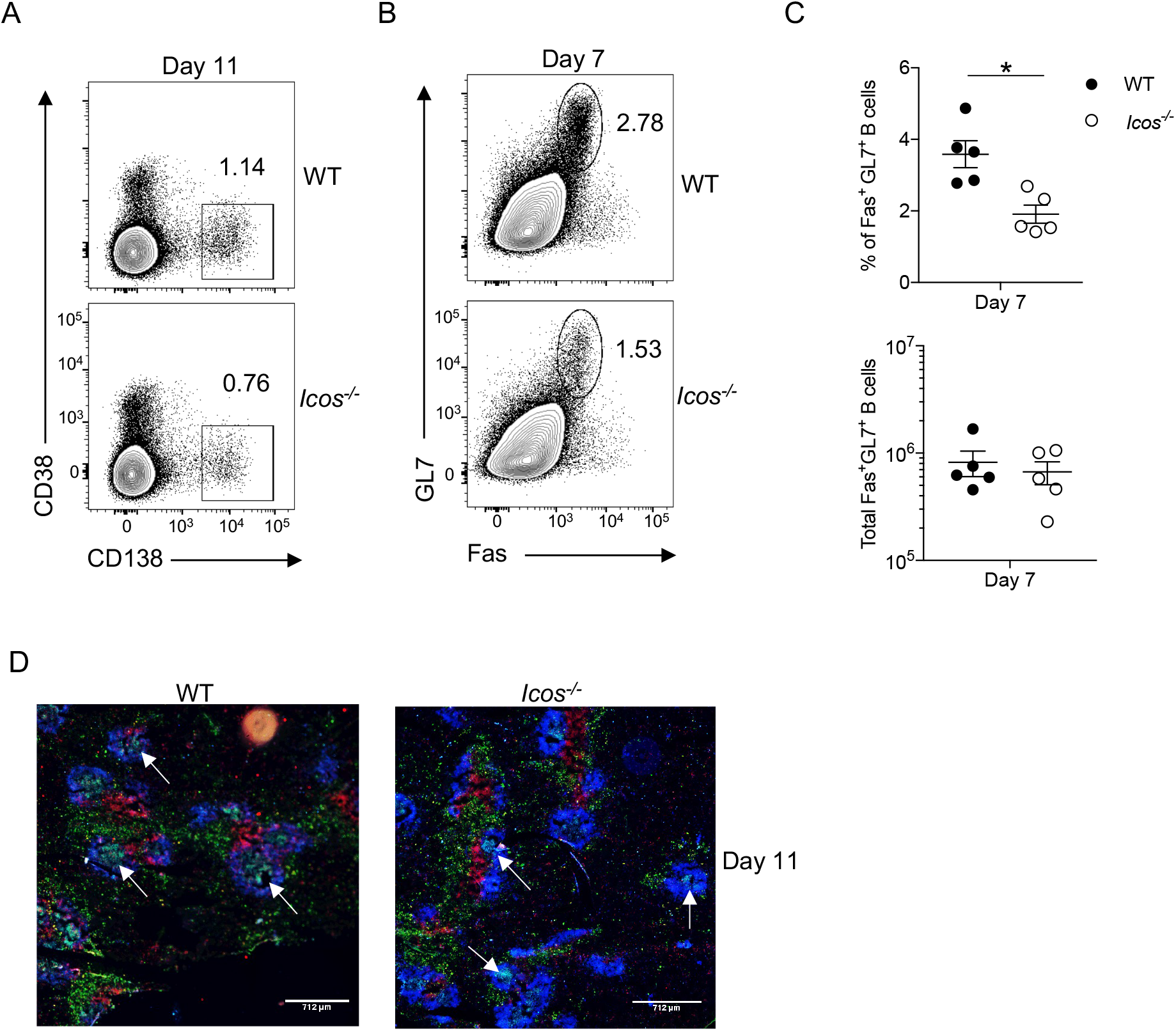
(A) Representative flow plots of CD138^+^CD38^-^ plasmablasts from WT and *Icos*^-/-^ mice at day 11 p.i. (B) Representative flow plots of Fas^+^GL7^+^ B cells at day 7 p.i. (**C**) Frequency and the total number of live CD19^+^B220^+^Fas^+^GL7^+^ B cells at day 7. (D) 4 μm spleen sections from WT and *Icos^-/-^* mice were stained with CD3*ε* (red), IgD (blue), and GL7 (green) at day 11 p.i. White arrows indicate representative GC structures. Data are representative of 3-4 independent experiments with five mice per group. (A, E, H, and I) An unpaired, two-tailed Student *t* test was used to determine significance. (B, D, and G) A nonparametric Mann-Whitney *t*-test determined significance. \**p* < 0.05.

**Supplemental FIG. 3.**
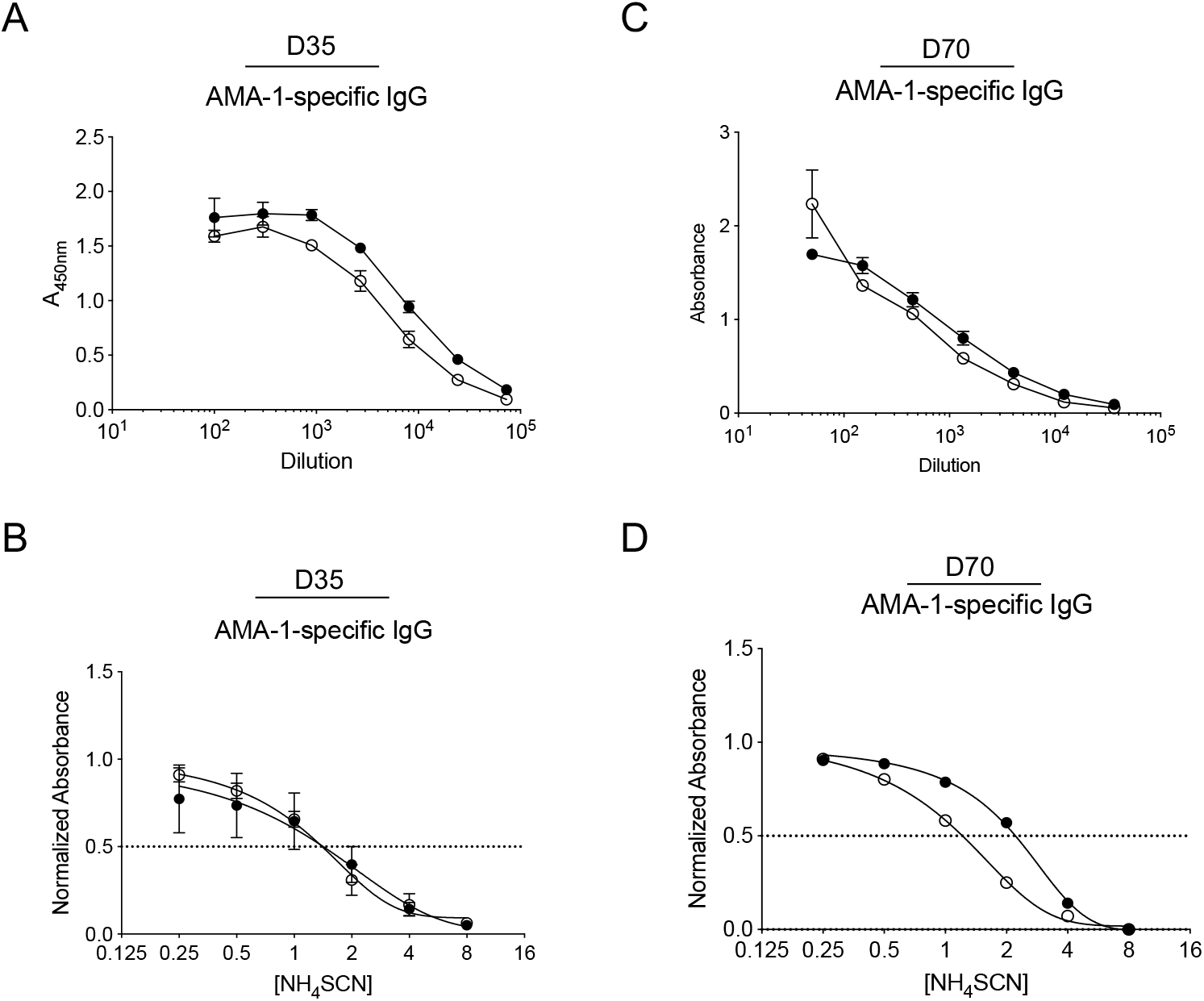
Serum AMA-1–specific IgG was determined by ELISA at **(A, B)** day 35 p.i. and **(C, D)** day 70 p.i. in WT and *Icos^-/-^* mice. Normalized AMA-1–specific IgG absorbance at **(B)** day 35 p.i. and **(D)** day 70 p.i. as determined by ELISA. Data are representative of two independent experiments with 4-5 mice per group. An unpaired, two-tailed Student *t* test determined significance.

**Supplemental FIG. 4.**
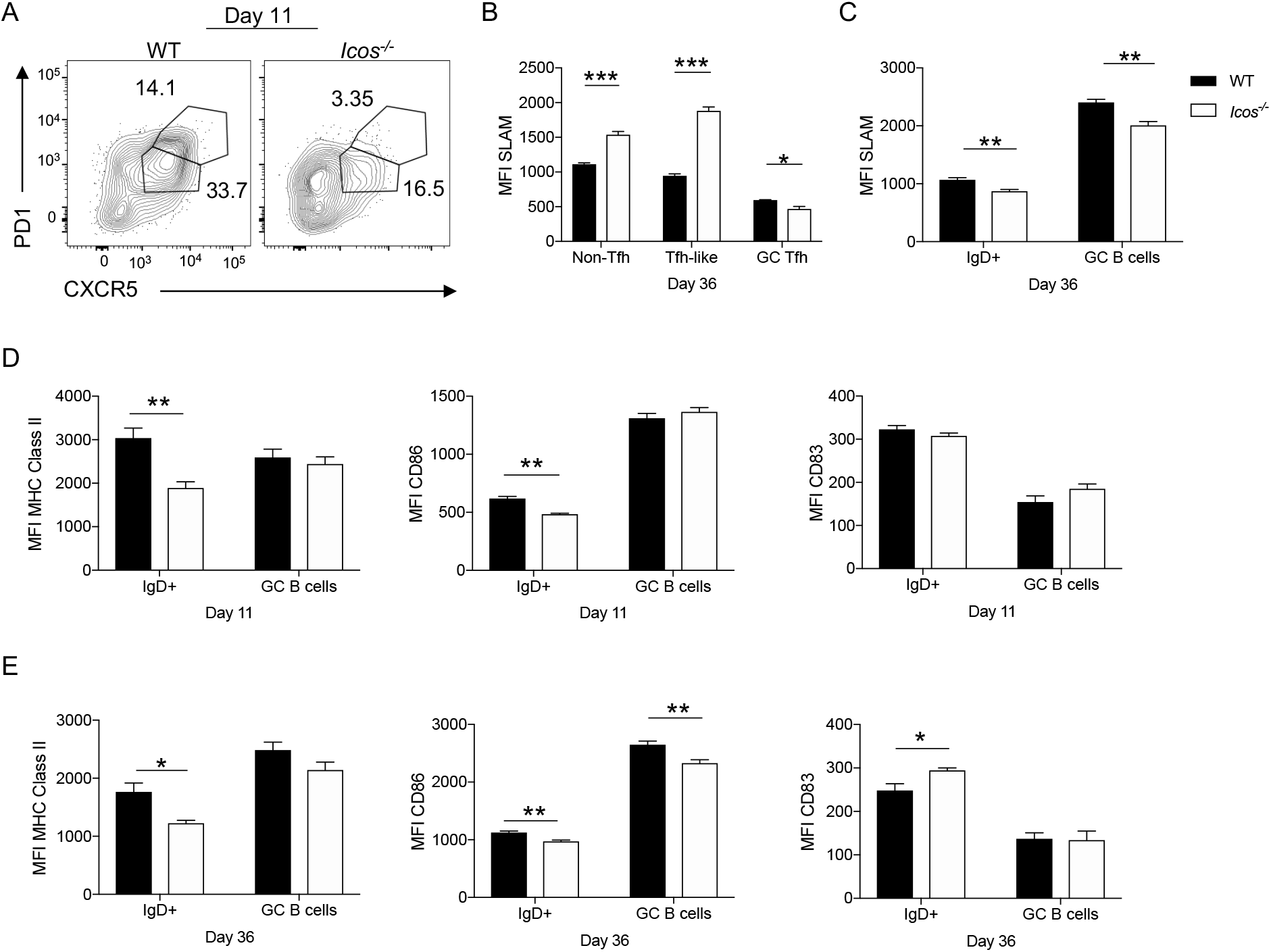
(A) Representative flow plot showing expression of CXCR5 and PD-1 on live activated CD44^hi^CD11a^+^ CD4^+^ T cells at day 11 p.i. **(B)** MFI of SLAM expression on live activated non-Tfh (CXCR5^-^PD-1^+/-^), Tfh-like, and GC Tfh CD4^+^ T cells at day 36 p.i. (**C**) MFI of SLAM expression on live CD19^+^B220^+^ IgD^+^ and Fas^+^GL7^+^ B cells at day 36 p.i. MFI of MHC class II, CD86, and CD83 expression on live CD19^+^B220^+^ IgD^+^ and Fas^+^GL7^+^ B cells at day (**D**) 11 and (**E**) 36 p.i. Data are representative of two independent experiments with five mice per group. Significance was determined by a nonparametric Mann-Whitney *t*-test. \**p* < 0.05, \*\**p* <0.01, \*\*\**p* < 0.001.

**Supplemental FIG. 5.**
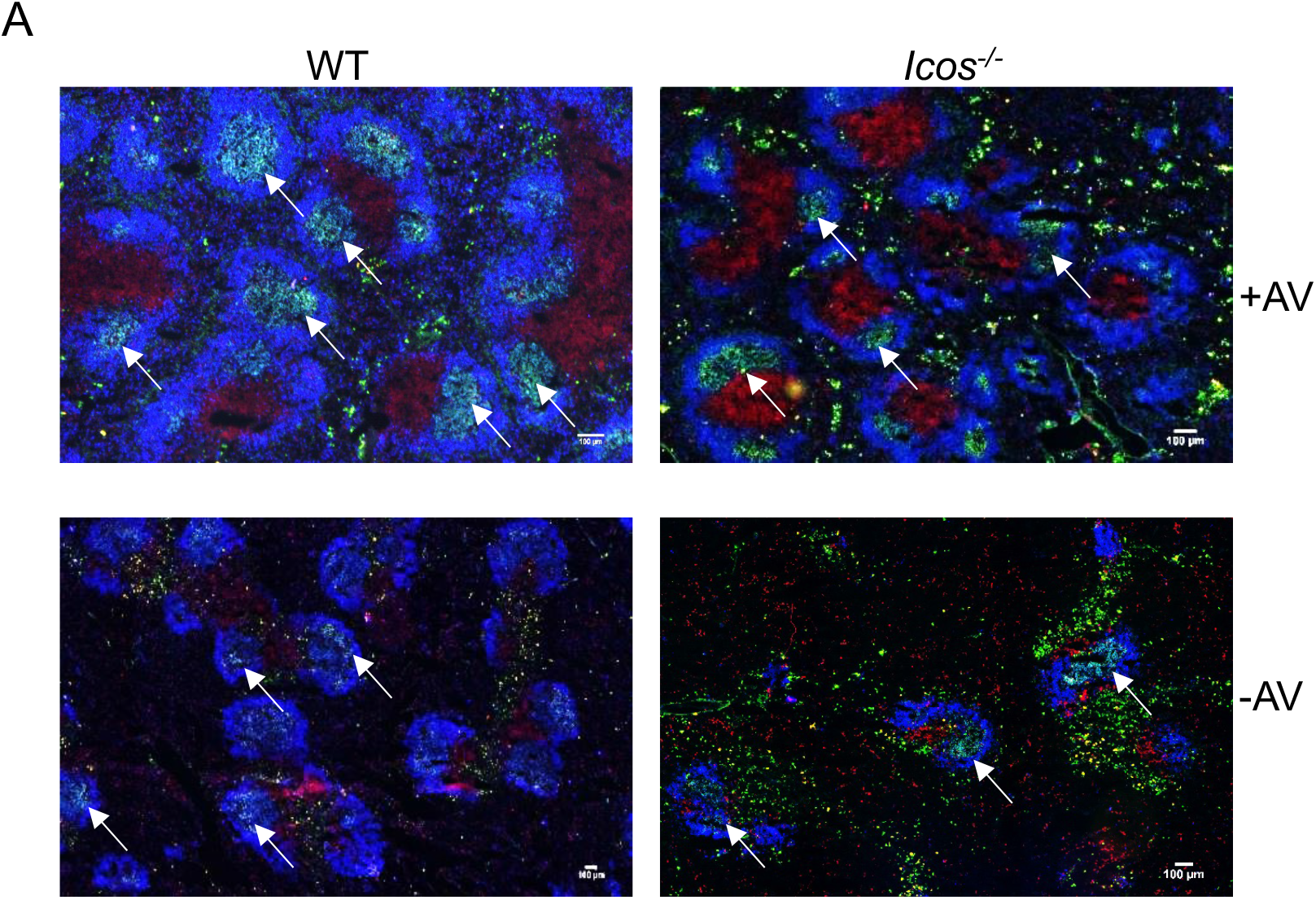
4 μm spleen sections from WT and *Icos^-/-^* mice were stained with CD3*ε* (red), IgD (blue), and GL7 (green) at day 21 p.i. White arrows denote representative GC structures. Scale bar is 100 μm. Data are representative of two independent experiments.

**Supplemental Table I.**
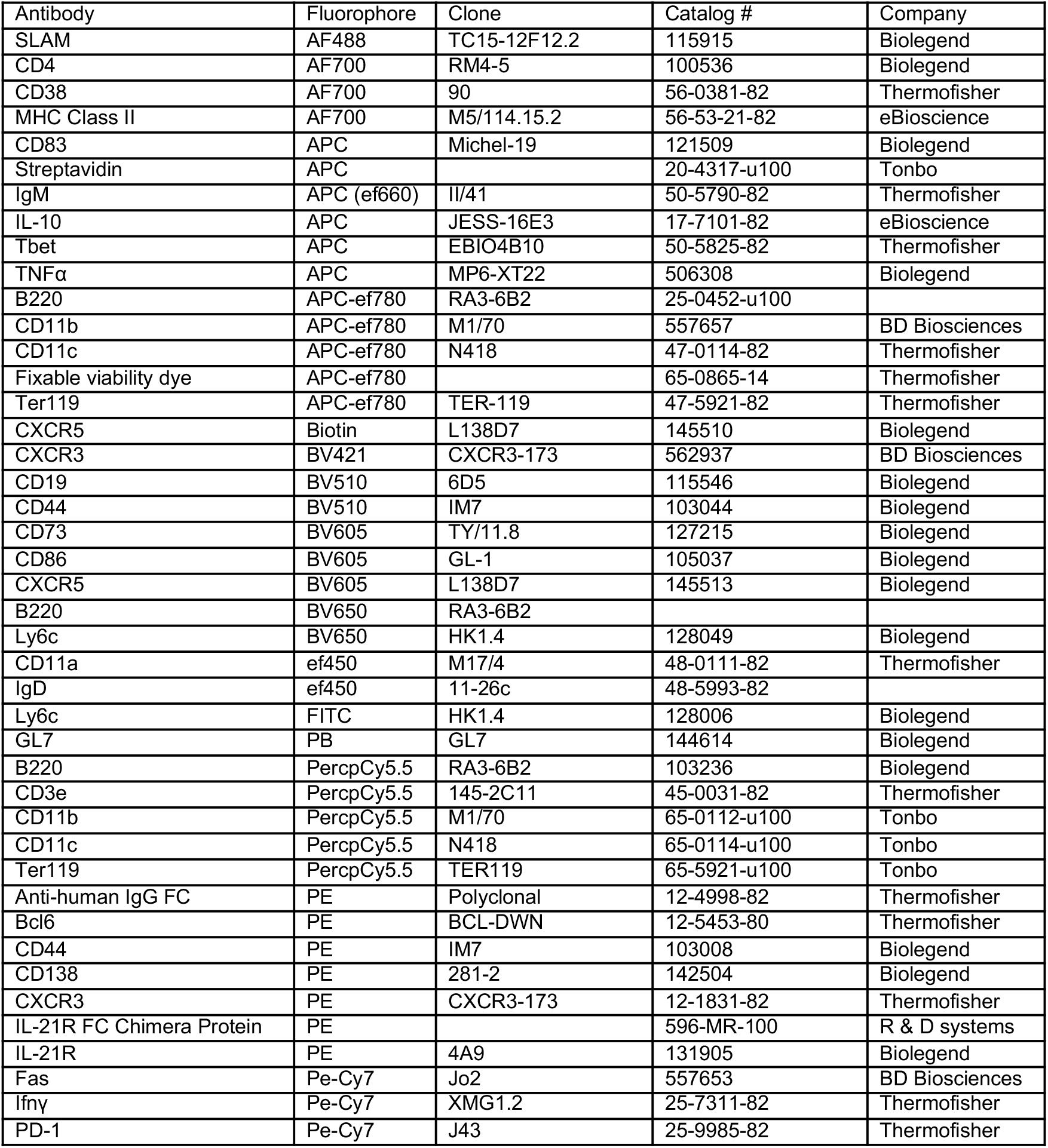
List of antibodies used for flow cytometry, including the fluorophore, clone, catalog number, and manufacturer.

**Supplemental Table II.**
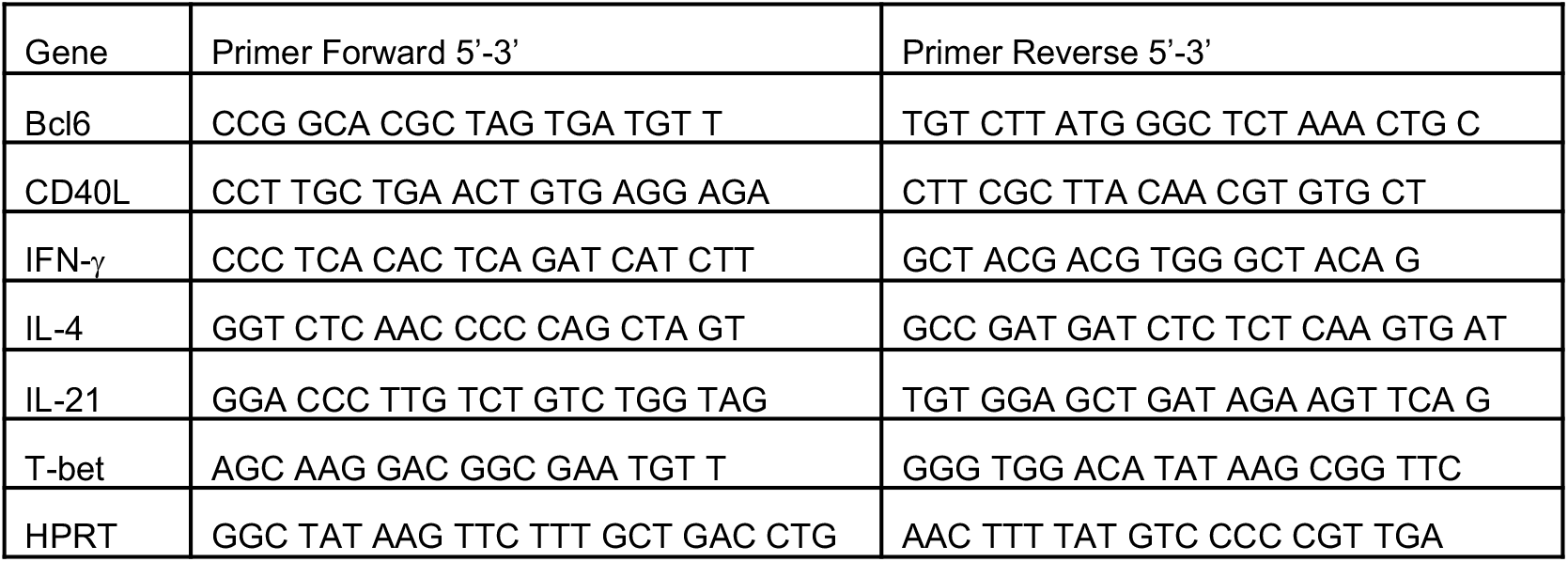
List of primer sequences used for real-time quantitative PCR analysis.

